# A single-cell atlas of lymphocyte adaptive immune repertoires and transcriptomes reveals age-related differences in convalescent COVID-19 patients

**DOI:** 10.1101/2021.02.12.430907

**Authors:** Florian Bieberich, Rodrigo Vazquez-Lombardi, Alexander Yermanos, Roy A. Ehling, Derek M. Mason, Bastian Wagner, Edo Kapetanovic, Raphael Brisset Di Roberto, Cédric R. Weber, Miodrag Savic, Fabian Rudolf, Sai T. Reddy

## Abstract

COVID-19 disease outcome is highly dependent on adaptive immunity from T and B lymphocytes, which play a critical role in the control, clearance and long-term protection against SARS-CoV-2. To date, there is limited knowledge on the composition of the T and B cell immune receptor repertoires [T cell receptors (TCRs) and B cell receptors (BCRs)] and transcriptomes in convalescent COVID-19 patients of different age groups. Here, we utilize single-cell sequencing (scSeq) of lymphocyte immune repertoires and transcriptomes to quantitatively profile the adaptive immune response in COVID-19 patients of varying age. We discovered highly expanded T and B cells in multiple patients, with the most expanded clonotypes coming from the effector CD8^+^ T cell population. Highly expanded CD8^+^ and CD4^+^ T cell clones show elevated markers of cytotoxicity (CD8: PRF1, GZMH, GNLY; CD4: GZMA), whereas clonally expanded B cells show markers of transition into the plasma cell state and activation across patients. By comparing young and old convalescent COVID-19 patients (mean ages = 31 and 66.8 years, respectively), we found that clonally expanded B cells in young patients were predominantly of the IgA isotype and their BCRs had incurred higher levels of somatic hypermutation than elderly patients. In conclusion, our scSeq analysis defines the adaptive immune repertoire and transcriptome in convalescent COVID-19 patients and shows important age-related differences implicated in immunity against SARS-CoV-2.

## Introduction

T and B lymphocytes are crucial for protection from SARS-CoV-2 infection, viral clearance and the formation of persisting antiviral immunity^1,2^. Yet, adaptive immune responses have also been implicated in contributing to immunopathology during COVID-19, with higher mortality rates in elderly individuals^3–6^. However, the exact determinants of a successful adaptive immune response against SARS-CoV-2 and its variability between different age groups remain to be fully elucidated.

Lymphocytes express either T cell receptors (TCR) or B cell receptors (BCR), which possess a highly diverse pair of variable chains [variable alpha (Vα) and beta (Vβ) for TCR and variable light (V_L_) and heavy (V_H_) for BCR] that are able to directly engage with antigen (e.g., viral proteins or peptides). Diversity in these variable chains are generated by somatic recombination of V-, D- and J-gene germline segments and along with combinatorial receptor chain pairing and somatic hypermutation (BCR only) results in an estimated human TCR and BCR diversity of 10^18^ and 10^13^, respectively^7,8^. Upon encountering cognate antigens, lymphocytes are phenotypically activated and undergo massive proliferation, also referred to as clonal selection and expansion. Deep sequencing of TCRs and BCRs has become a powerful strategy to profile the diversity of immune repertoires and to reveal insights on clonal selection, expansion and evolution (somatic hypermutation in B cells)^9–12^ and has been instrumental in studying long term effects following vaccination, infection and ageing^13–17^. In the context of COVID-19, immune repertoire sequencing has shown diminished TCR repertoire diversity and BCR isotype switching and respective expansion during early disease onset^18^.

In recent years, single-cell sequencing (scSeq) of transcriptomes has progressed substantially through the development and integration of technologies such as cell sorting, microwells and droplet microfluidics^19,20^; most notably commercial systems like those of 10X Genomics have been established and are providing standardized protocols for wider implementation of scSeq. To find interactions across multiple genes and cells, analysis and visualisation of this high dimensional single-cell data is facilitated by clustering and nonlinear dimensionality reduction algorithms [e.g., t-distributed stochastic neighbor embedding (t-SNE) or Uniform Manifold Approximation and Projection (UMAP)]^21,22^. scSeq of transcriptomes has been used extensively to profile the gene expression signatures of T and B cells to identify novel cellular subsets and phenotypes as well as their response to vaccination, infection and cancer^23–26^. Furthermore, clustering with scSeq data enables the unbiased identification of cellular states and analyses of the broad continuum of T and B cell populations as well as their differentiation trajectories^27^. In the context of patients with severe symptoms of COVID-19, scSeq has revealed a dysfunctional T cell response of interferon expression combined with elevated levels of exhaustion^28^.

In addition to transcriptome sequencing, a major advantage of scSeq is that it also enables information on the native pairing of TCR Vα and Vβ chains and BCR V_L_ and V_H_ chains^29–32^, which was not previously possible with the standard bulk sequencing of lymphocytes as these receptor chains are expressed as unique transcripts from separate chromosomes ^33^. Coupling TCR or BCR sequence to the transcriptome within an individual cell enables phenotypic analyses of a clonal population of lymphocytes and their dynamics^34–36^. scSeq of transcriptomes and immune repertoires in COVID-19 patients with severe symptoms has shown a high level of clonal expansion in specific T cell subsets (Th1, Th2, and Th17) and preferential germline gene usage in clonally expanded B cells^28,34,37^; while a more recent study found a positive correlation between clonal expansion of effector-like CD8+ T cells and disease severity^38^.

An important question that remains to be answered is whether there are age-related differences in mounting a successful adaptive immune response against SARS-CoV-2. Here, we perform scSeq on the immune repertoires and transcriptomes of T and B cells derived from eight convalescent COVID-19 patients of two different age groups (mean ages = 31 and 66.8 years) at one month of convalescence following mild to moderate disease. We observed preferential clonal expansion of effector CD8+ T cells across all patients, although a significantly higher CD8-to-CD4 T cell ratio was detected in young patients of our cohort. Further, clonally expanded B cells in young patients displayed significantly higher levels of somatic hypermutation and an increased immunoglobulin (Ig) class-switching compared to clonally expanded B cells from older patients. Our analyses serve as a valuable resource for future scSeq characterization of SARS-CoV-2 adaptive immunity and highlight important age-related differences in the adaptive immune status of convalescent COVID-19 patients.

## Results

### Study design and single-cell profiling of convalescent COVID-19 patient lymphocytes

We performed scSeq of immune receptor repertoires and transcriptomes of lymphocytes from convalescent COVID-19 patients to characterize the adaptive immune response against SARS-CoV-2. For this purpose, we selected eight patients enrolled in the SERO-BL-COVID-19 clinical study^39^, all of which fully recovered from COVID-19 without requiring hospitalization or the administration of Supplementary oxygen. Patients tested positive for the presence of SARS-CoV-2 after RT-PCR of naso/oropharyngeal swab samples (day 0), displayed COVID-19 symptoms for 4-14 days, and showed positive seroconversion at the time of blood collection (mean sample collection time = 32.5 ± 4.1 days post-symptom onset) (**Fig. 1a and Supplementary Table 1**). Since COVID-19 often affects older patients more severely^40^, subjects were divided into two groups according to their age, namely Group 1 (mean = 66.75 ± 6.9 years) and Group 2 (mean = 31 ± 5.9 years), with the aim of investigating potential differences in their responses against SARS-CoV-2. In addition to older age, significant differences in Group 1 versus Group 2 included elevated IgA/IgG SARS-CoV-2-specific antibody titers and an increased duration of COVID-19 symptoms (**Fig. 1b and Supplementary Fig.1**). Patients from Group 1 also experienced an increased severity of COVID-19 symptoms relative to Group 2 (**Supplementary Table 1**). Despite increased symptom duration in the older cohort, correlation of this parameter with age was only modest (R^2^ = 0.4647), likely reflecting the small sample size (**Supplementary Fig. 1c**).

**Figure 1.**
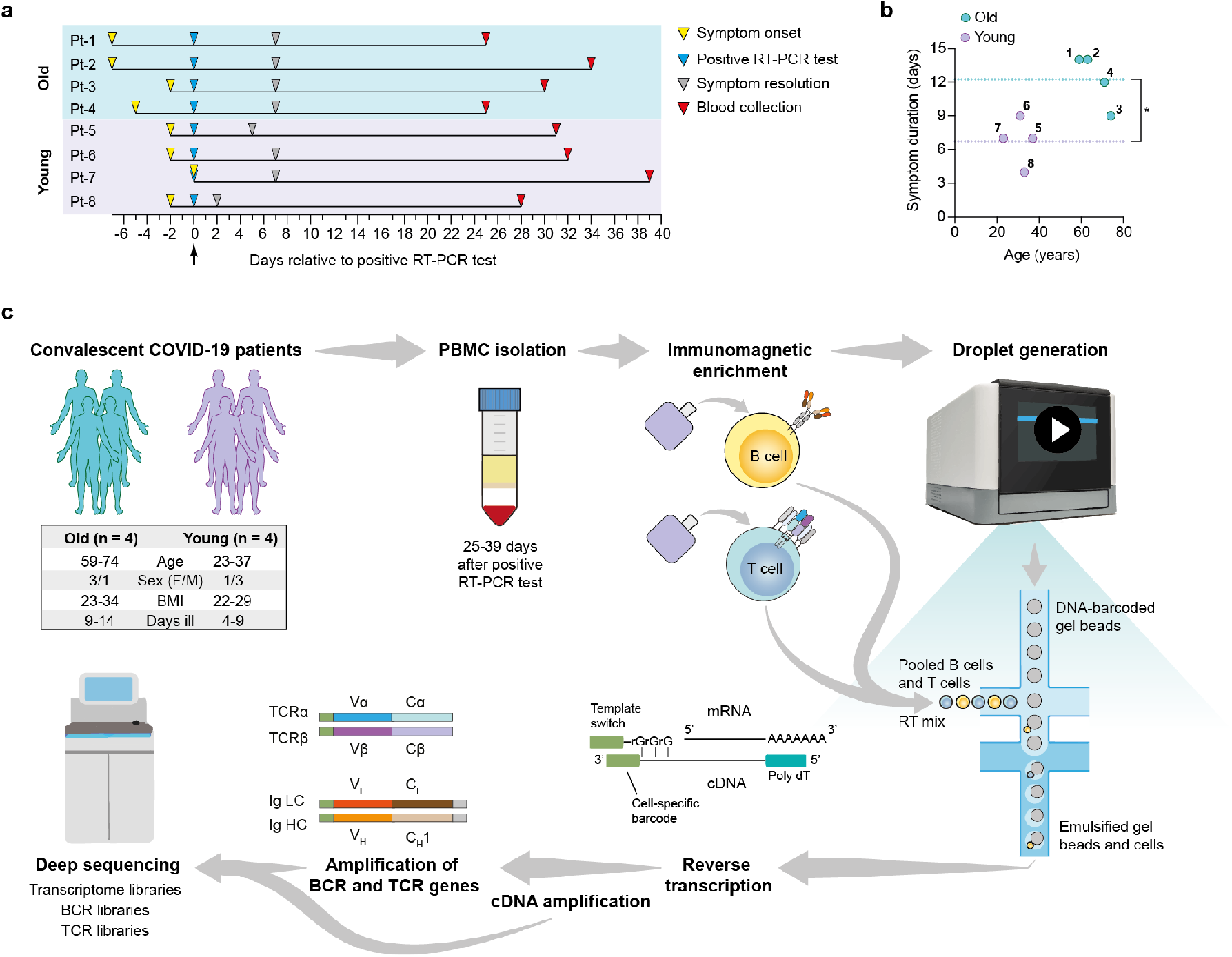
Overview of single-cell transcriptome and immune receptor profiling of convalescent COVID-19 patient lymphocytes. Convalescent COVID-19 patients enrolled in the SERO-BL-COVID-19 study were selected according to their age for single-cell sequencing analysis of their T cells and B cells. **a**, Timeline illustrates symptom onset, symptom resolution and collection of blood samples from individual patients relative to the time of positive SARS-CoV-2 RT-PCR test (day 0). **b**, Graph displays the ages and duration of COVID-19 symptoms in individual patients. Dotted lines show the mean duration of symptoms in the young (y = 6.75 days) and old (y = 12.25 days) groups. A significant difference in symptom duration between groups is indicated with an asterisk (p = 0.0127; unpaired t-test). **c**, Single-cell sequencing protocol. Whole blood was collected following the resolution of COVID-19 symptoms and subjected to density gradient separation for isolation of PBMC. T cells and B cells from individual patients were purified from PBMC using negative immunomagnetic enrichment, pooled (intra-patient) and prepared for droplet generation using the 10x Genomics Chromium system. Single cells were emulsified with DNA-barcoded gel beads and mRNA transcripts were reverse-transcribed within droplets, resulting in the generation of first-strand cDNA molecules labelled with cell-specific barcodes at their 3’ ends (added by template switching). Emulsions were disrupted and cDNA was amplified by means of PCR for further processing of transcriptome libraries. Transcriptome libraries from individual patients were indexed and multiplexed for deep sequencing using the Illumina NovaSeq platform. Targeted enrichment of recombined V(D)J transcripts was performed by PCR and the resulting products were processed for the generation of BCR and TCR libraries, which were then indexed, multiplexed and deep-sequenced.

To profile patient lymphocytes, we isolated peripheral blood mononuclear lymphocytes (PBMC) from blood and purified T cells and B cells by negative immunomagnetic enrichment. Plasma cells (PCs) were depleted from PBMC samples prior to this step for scSeq in a companion study (Ehling et al., manuscript in preparation), and thus were excluded from our analyses. After purification, T cells and B cells underwent the 10X genomics protocol for scSeq 5’ library preparation, which included gel encapsulation single-cell barcoding of mRNA, followed by cDNA generation through polydT reverse transcription. Finally, after full-length V(D)J segment enrichment, construction of TCR and BCR V(D)J and transcriptome sequencing libraries was done according to the V(D)J enrichment and 5’ library construction kits, respectively. Deep sequencing of immune repertoires and transcriptomes was performed using the Illumina NovaSeq with paired-end 26 x 91 bp cycles per read. For TCR and BCR V(D)J and transcriptome libraries, we recovered on average 20.000 and 10.000 reads per cell, respectively (**Fig. 1c**).

### Single-cell transcriptome analysis defines major T and B cell subsets

Bioinformatic filtering was performed to exclude the following: Cell doublets, cells with a very low or high number of genes, and T cells with no detectable expression of CD8 and CD4 (see Methods), which resulted in the identification of 30,096 cells in total from all eight patients. Cells were then split into CD8+ T cell (**Fig. 2a**, 7,353 cells), CD4+ T cell (**Fig. 2b**, 8,334 cells) and B cell (**Fig. 2c** 14,409 cells) datasets. In order to reduce the dimensionality of the data, while preserving the global structure, we used UMAP for better visualisation and interpretation purposes^41^. UMAP and unsupervised clustering of these subgroups led to the identification of eleven dominant cell subsets (**Figs. 2a-c**). CD8+ T cells clustered into naïve (SELL+, TCF7+), memory (IL7R+. CD40LG+) and effector cells (GZMB+, NKG7+) (**Figs. 2a and 2d**), which also encompassed exhausted CD8+ T cells (**Supplementary Fig. 2**). We identified four different CD4+ T cell subsets, namely naïve (SELL+, LEF1+), memory (S100A4+), effector (CCL5+, GZMK+) and regulatory cells (FOXP3+) (**Figs. 2b and 2e**). The B cell compartment consisted of naïve (CD23A+) marginal zone (MZ) (FCRL3+, CD1C+), activated (CD83+) and memory cells (CD27+, TACI+) (**Figs. 2c and 2f**). Notably, clustering of single cells based on transcriptome data revealed a trajectory that reflected a progression in lymphocyte differentiation from naïve to effector (or activated) subsets (**Figs. 2a-c**). Pseudotime analysis of the dataset supports this differentiation trajectory in CD8+ and CD4+ T cells (**Figs. 2g and 2h**). Interestingly, pseudotime analysis of B cell data not only showed naïve-MZ-memory and naïve-MZ-activated trajectories, but also a third MZ-memory-activated trajectory that suggests the presence of reactivated memory B cells, possibly through antigen encounter (**Fig. 2i**).

**Figure 2.**
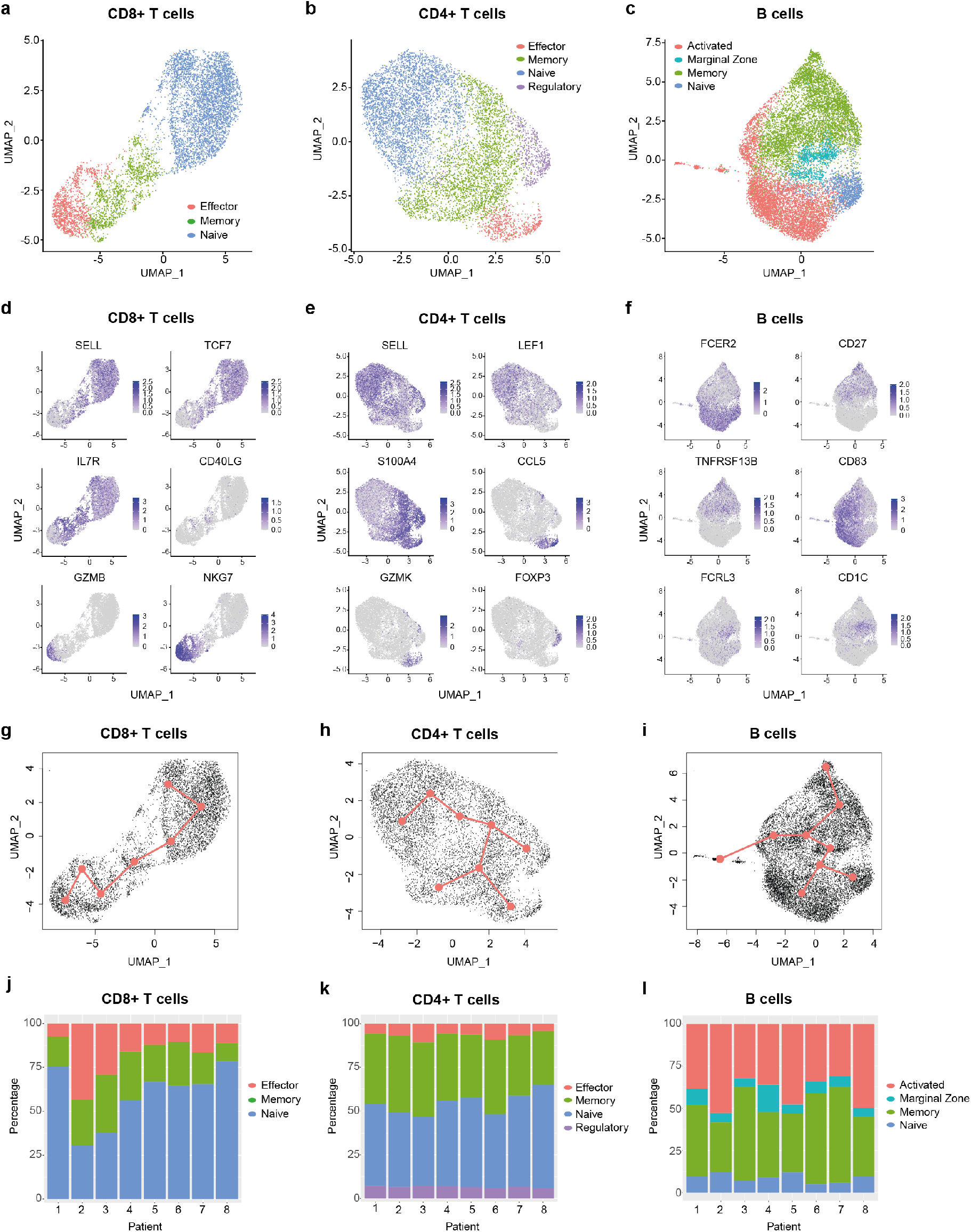
Single-cell transcriptomic analysis delineates major T and B cell subsets. **a-c**, Uniform manifold approximation and projection (UMAP) plots of major cellular subsets identified within the CD8+ T cell (**a**), CD4+ T cell (**b**) and B cell (**c**) populations. Cells from all patients are displayed in each plot. **d-f**, UMAP plots showing the expression levels of selected genes used to delineate cellular subsets within the CD8+ T cell (**d**), CD4+ T cell (**e**) and B cell (**f**) populations. Cells from all patients are displayed in each plot. **g-i**, Graphs display pseudotime and trajectory inference analysis applied to CD8+T cell (**g**), CD4+ T cell (**h**) and B cell (**i**) clusters. **j-l**, Bar graphs show the proportions of identified cellular subsets within the CD8+ T cell (**j**), CD4+ T cell (**k**) and B cell (**l**) populations in each patient. CCL5 = C-C Motif Chemokine Ligand 5; CD27 = TNFRSF7; CD40LG = CD40 ligand; FCER2 = Fc Fragment of IgE Receptor II (also: CD23a); FCRL3 = Fc Receptor Like 3; FOXP3 = Forkhead Box Protein P3; GZMB = Granzyme B; GZMK = Granzyme K; IL7R = Interleukin-7 Receptor; LEF1 = Lymphoid Enhancer Binding Factor 1; NKG7 = Natural Killer Cell Granule Protein 7; S100A4 = S100 Calcium Binding Protein A4; SELL = Selectin L; TCF7 = Transcription Factor 7; TNFRSF13B = TNF Receptor Superfamily Member 13B.

Having defined the major T cell and B cell subsets from pooled patient data, we next compared their proportions across patients (**Figs. 2j-l**) and between different age groups (**Supplementary Fig. 3**). We found that young patients had a significantly higher CD8-to-CD4 T cell ratio relative to older ones (**Supplementary Fig. 3a**), which may reflect a previously reported age-dependent difference^42^. Interestingly, despite this reduction, there was a trend that older patients had a higher proportion of effector CD8+ T cells relative to their younger counterparts (**Supplementary Fig. 3b**). While this difference was not significant, it is consistent with the increased symptom severity experienced by older patients (**Supplementary Table 1**), a feature that has been associated with elevated proportions of effector/exhausted CD8+ T cells in the periphery^28^. Of note, we found that older patients had a small but significant increase in CD4+ Tregs compared to young patients (**Supplementary Fig. 3h**), and that increased proportions of MZ B cells occurred in two of the older patients (**Supplementary Fig. 3l**). Taken together, our data highlights the diversity of elevated responses in specific patients across age groups, as exemplified by individuals with a high abundance of effector CD8+ T cells (e.g., Pt-2 and Pt-3) and/or activated B cells (e.g., Pt-2, Pt-5 and Pt-8).

### Single-cell profiling of immune receptor repertoires identifies highly expanded TCR and BCR clonotypes

We next determined the clonal expansion levels of T cells and B cells in convalescent COVID-19 patients by quantifying the number of cells expressing unique TCRs (**Fig. 3a**) or BCRs (**Fig. 3c**) (clonotype definition in the methods section). We found substantial heterogeneity in T cell clonal expansion levels across patients, with the highest number of expanded TCR clonotypes occurring in four patients, namely Pt-2 and Pt-3 (Group 1), Pt-7 and Pt-8 (Group 2). Within these patients, Pt-2 displayed the largest amount of expanded TCR clonotypes, which is consistent with the high abundance of effector CD8+ T cells in this subject (**Fig. 2g**). Analysis of TCRα and TCRβ germline V-gene usage in the ten most expanded clonotypes per patient revealed a frequent occurrence of TRBV20-1 (7 out of 8 patients) and TRAV-29/DV5 genes (5 out of 8 patients), though pairing of these germline genes was not observed (**Fig. 3b and Supplementary Fig. 4**). In agreement with the overall higher expansion of CD8+ effector over CD4+ effector T cell subsets (**Figs. 2g and 2h**), we found that the vast majority (85%) of the ten most expanded TCR clonotypes per patient originated from CD8+ T cells (**Supplementary Fig. 5**). Based on this observation, we genotyped patient HLA class I alleles by means of amplicon deep sequencing (**Supplementary Table 2**). We found that the two patients with the highest levels of T cell clonal expansion (i.e., Pt-2 and Pt-8) shared the HLA-A*0201 allele, as well as a number of TRBV and TRAV genes in their most expanded clonotypes, which indicates a possible convergence towards germlines that may be related to SARS-CoV-2 specificity. Analysis of single-cell BCR repertoire sequencing data revealed that highly expanded BCR clonotypes occurred more frequently in younger patients, for example in Pt-5, Pt-6 and Pt-8 (**Fig. 3c**). This is an unexpected finding, particularly as older patients in our cohort displayed significantly higher SARS-CoV-2-specific IgA/IgG titers in serum (**Supplementary Figs. 1e and 1f**). Thus, this suggests that older patients may harbor a higher diversity of relatively unexpanded SARS-CoV-2-specific B cells. Supporting this observation, we found that older patients had a wider range of heavy chain complementarity determining region 3 (CDR3H) lengths relative to younger ones, indicating a possible larger degree of variability in their antibody paratopes (**Supplementary Fig. 6**). Analysis of heavy chain and light chain germline V-gene pairing in the ten most expanded BCR clonotypes per patient revealed a frequent occurrence of the IGHV-3-23 / IGKV-3-20 pairing (7 out 8 patients) (**Fig. 3d and Supplementary Fig. 7**). However, as this pairing is the most frequently found in healthy cohorts^43,44^, such antibodies may not necessarily be enriched for SARS-CoV-2 specificity.

**Figure 3.**
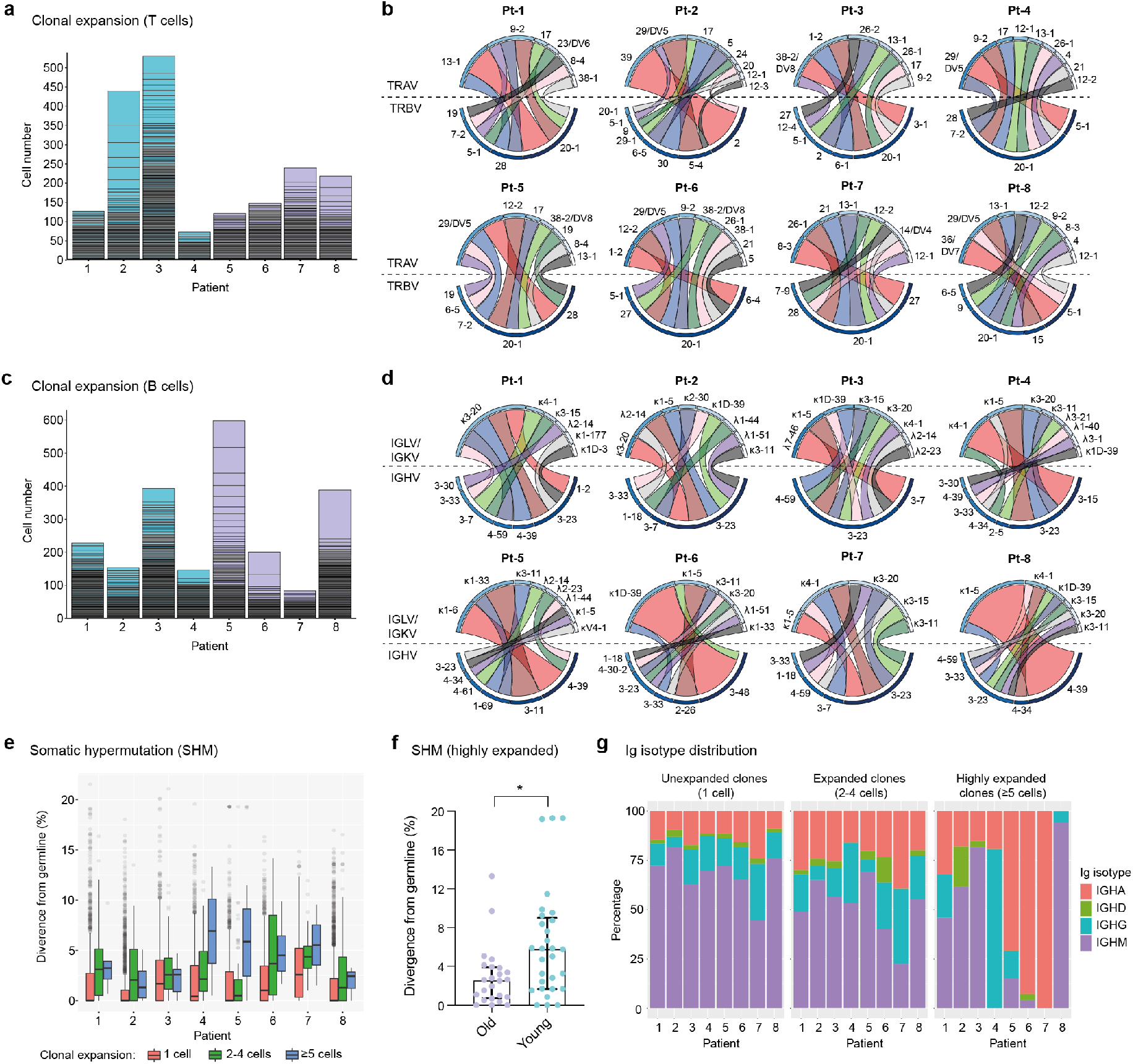
Single-cell profiling of immune repertoires highlights differential levels of inter-patient T cell and B cell clonal expansion. **a-b**, Analysis of T cell clonal expansion in convalescent COVID-19 patients. **a**, Bar graphs show T cell clonal expansion, as determined by the number of cells identified per TCR clonotype. Each box represents the size of individual TCR clonotypes. TCR clonotypes present in more than one cell are shown. **b**, Circos plots display V-gene usage in the top ten most expanded TCR clonotypes for each patient. The size and colour (dark to light) of outer bars reflect the relative abundance of T cells expressing specific V-genes on a per class basis (top: TCRα chain, bottom: TCRβ chain). **c-e**, Analysis of B cell clonal expansion in convalescent COVID-19 patients. **c**, Bar graphs show B cell clonal expansion, as determined by the number of cells identified per BCR clonotype. Each box represents the size of individual BCR clonotypes. BCR clonotypes present in more than one cell are shown. **d**, Circos plots display V-gene usage in the top ten most expanded BCR clonotypes for each patient. The size and colour (dark to light) of outer bars reflect the relative abundance of B cells expressing specific V-genes on a per class basis (top: Ig light chain, bottom: Ig heavy chain). **e**, Graph displays the levels of somatic hypermutation (SHM) in unexpanded (1 cell), expanded (2-4 cells) and highly expanded (5 cells) BCR clonotypes across patients. SHM levels are based on the percentage similarity between Ig heavy chain V-genes and their corresponding germlines. Data are displayed as median ± IQR. **f**, Graph displays SHM levels in highly expanded BCR clones (≥5 cells) of old (n = 24 clones) and young (n = 29 clones) patients. Asterisks indicate a significant difference in SHM levels between groups (p = 0.0085; unpaired t-test). Data are displayed as median ± IQR. **g**, Bar graphs show the Ig isotype distribution in unexpanded (1 cell), expanded (2-4 cells) and highly expanded (≥5 cells) BCR clonotypes across patients.

To further characterize BCR repertoires across different levels of clonal expansion we divided clonotypes into additional subsets: unexpanded (1 cell per clonotype), expanded (2-4 cells per clonotype) and highly expanded (≥5 cells per clonotype) and assessed their levels of somatic hypermutation (SHM). We found that the degree of SHM largely correlated with clonal expansion, with expanded and highly expanded clonotypes having higher SHM (i.e., more divergent from their germline V-genes) than unexpanded ones (**Fig. 3e**). Strikingly, highly expanded BCR clonotypes from young patients had significantly higher SHM levels compared to older patients, potentially indicating more efficient affinity maturation had occurred in response to SARS-CoV-2 antigens (**Fig. 3f**). Finally, we examined the distribution of Ig isotypes across clonal expansion groups. As expected, IgM was the most frequent isotype in unexpanded BCR clonotypes, with the proportion of this isotype being reduced in expanded BCR clonotypes (2-4 cells) of all patients. Conversely, the proportions of IgG and IgA isotypes in expanded clonotypes increased for all patients, thus indicating class-switching in response to clonal expansion. Analysis of Ig isotype distribution in highly expanded BCR clonotypes (≥5 cells) revealed that a subset of patients harbored a vast majority of class-switched IgA (Pt-5, Pt-6 and Pt-7) or IgG (Pt-4). Notably, we found that Ig isotype class-switching in highly expanded clonotypes was correlated with SHM levels across patients (Fig. 3e), highlighting the temporal connection between affinity maturation and class-switching processes in the germinal center^45^.

### Single-cell transcriptome and TCR profiling reveals predominant cytotoxic programs in highly clonally expanded CD4+ and CD8+ T cells

We next investigated the patterns of clonal expansion in different T cell subsets by mapping single-cell TCR sequencing data onto individual CD8+ and CD4+ T cells visualized by UMAP (**Figs. 4a and 4b**). For this analysis, we identified a total of 4,730 CD8+ T cells and 5,509 CD4+ T cells with available TCR clonotype and transcriptome information. Both CD8+ and CD4+ T cells showed increased levels of clonal expansion when progressing from naïve to effector phenotypes, with highly expanded TCR clonotypes (≥5 cells) almost exclusively expressed by effector T cells (**Figs. 4a and 4b, Figs. 2a and 2b**). As previously observed (**Supplementary Figs. 3 and 5**), CD8+ T cells showed substantially higher levels of clonal expansion relative to CD4+ T cells, in which highly expanded clonotypes were rare. Notably, young patients (Group 2) had a markedly higher abundance of unexpanded CD8+ T cell clonotypes compared to older patients (Group 1), which could indicate an ongoing resolution of their CD8+ T cell response at the analyzed timepoint. Patients with high levels of CD8+ T cell clonal expansion, however, were identified across age groups (i.e., Pt-2, Pt-3, Pt-7 and Pt-8), with Pt-2 (Group 1) showing the highest abundance of highly expanded clonal T cells. We further explored the relationship between clonal expansion and T cell phenotype by performing differential gene expression analysis in unexpanded, expanded and highly expanded T cell clonotypes (**Figs. 4c and 4d**). CD8+ T cells with high clonal expansion displayed elevated cytotoxicity (PRF1, GZMH, GNLY), activation (NKG7, CCL5), inflammation (NFGBIA, S100A4, S100A6) and type I interferon-induced (IFITM2) markers in all patients. Additionally, components of MHC class I (HLA-A, HLA-B, HLA-C and B2M) were also increased in this subgroup, indicating increased IFN-γ-induced activation^46^. Conversely, unexpanded CD8+ T cell clonotypes across age groups displayed upregulated markers found in naïve and memory CD8+ T cell subsets (IL7R, LTB)^47^, as well as markers likely associated with homeostatic proliferation (LDHB, NOSIP, EEF1B2, NPM1, TPT1, PABPC1).

**Figure 4.**
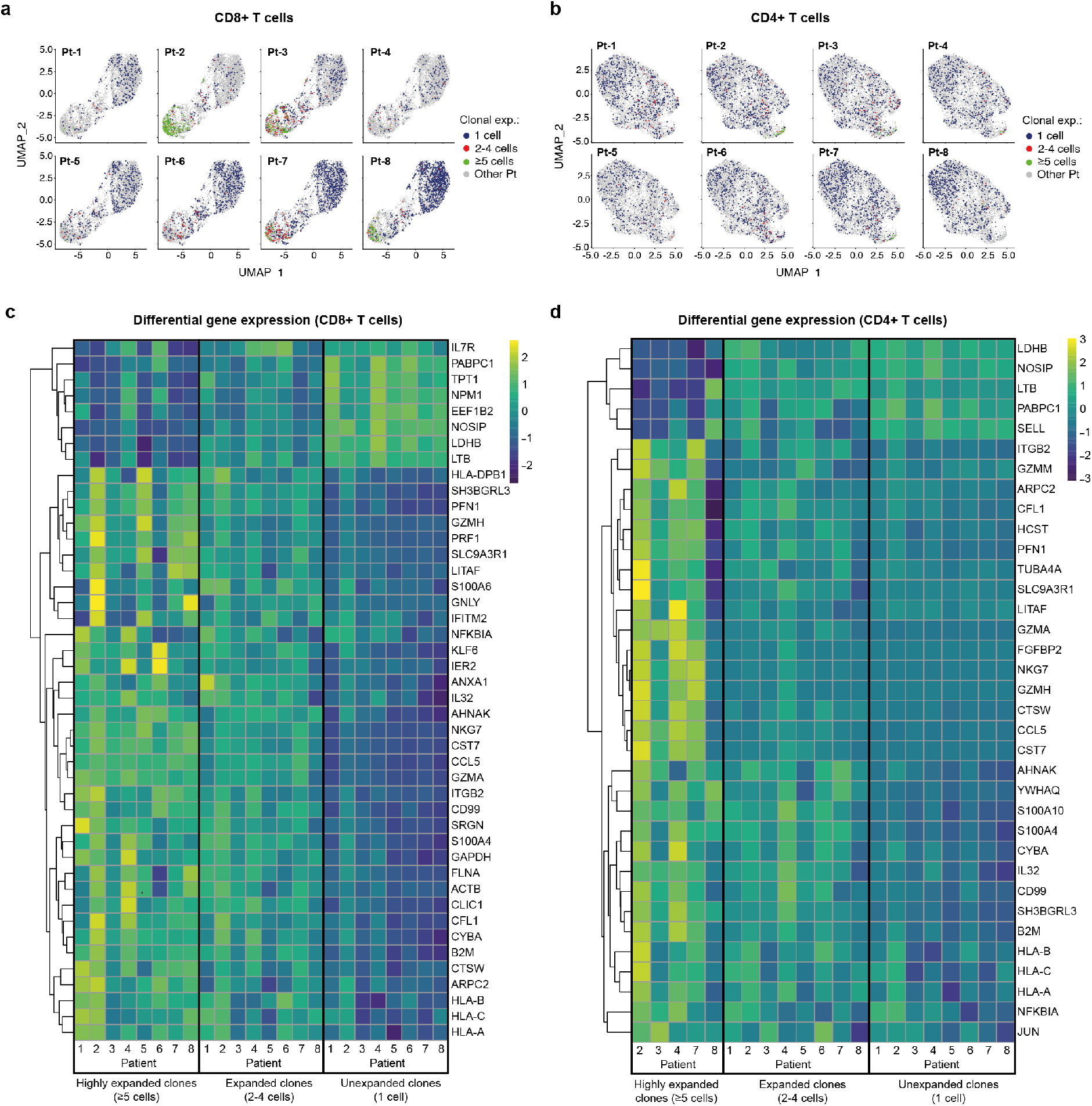
Single-cell transcriptome and TCR sequencing reveals preferential clonal expansion in effector T cells. **a-b**, UMAP plots display CD8+ (**a**) and CD4+ (**b**) T cells from specific patients according to their clonal expansion levels. T cells from other patients in each individual plot are shown in grey. **c-d**, Heatmaps show differential gene expression (DGE) in unexpanded, expanded and highly expanded CD8+ (**c**) or CD4+ (**d**) T cells. Genes were filtered to include those with detectable expression in at least 50% of cells and that had a minimum 50% fold-change in expression level between groups.

Despite the low levels of clonal expansion observed in CD4+ T cells, we identified distinct gene expression signatures in the highly expanded clonotypes that were present in 5 of 8 patients. Similar to CD8+ T cells, highly expanded CD4+ T cell clonotypes showed upregulation of genes related to activation (CCL5), cytotoxicity (GZMA), inflammation (IL32, CD99, NFKBIA) and MHC class I molecules (HLA-A, HLA-B, HLA-C and B2M), while unexpanded CD4+ T cells displayed markers of naïve T cells (SELL) and proliferation markers (LDHB, NOSIP, PABPC1). Taken together, integration of TCR sequencing and transcriptome data reveals clonal expansion as a hallmark of effector T cell subsets, and highlights a dominant role of CD8+ T cells in possible clonal responses against SARS-CoV-2 in convalescent COVID-19 patients.

### Transcriptomic and BCR profiling of single B cells reveals plasma cell transition, class-switching and SHM patterns

We next integrated single-cell BCR sequencing data onto the B cell transcriptional landscape to relate clonal expansion, SHM and isotype distribution to different B cell phenotypes. For this analysis we identified 11,227 individual B cells with available BCR and transcriptomic information. We observed a generally low level of B cell clonal expansion, with preferential localization of expanded B cell clonotypes (2-4 cells) to the memory and MZ B cell regions and a rare occurrence of highly expanded (≥5 cells) clonotypes in most patients (**Fig. 5a**). Analysis of differential gene expression showed that expanded clonotypes had increased expression of genes involved in cytoskeleton reorganization (VIM) and genes associated with the unfolded protein response (HSBP90, CALR, PPIB), indicating B cell activation and transition into plasma cells, respectively^48–50^ (**Fig. 5b**). Furthermore, expanded B cell clonotypes across patients showed downregulation of MHC class II genes (CD74, HLA-DR, -DQA1, -DRB1), further supporting their trajectory towards antibody-producing plasma cells^51^.

**Figure 5.**
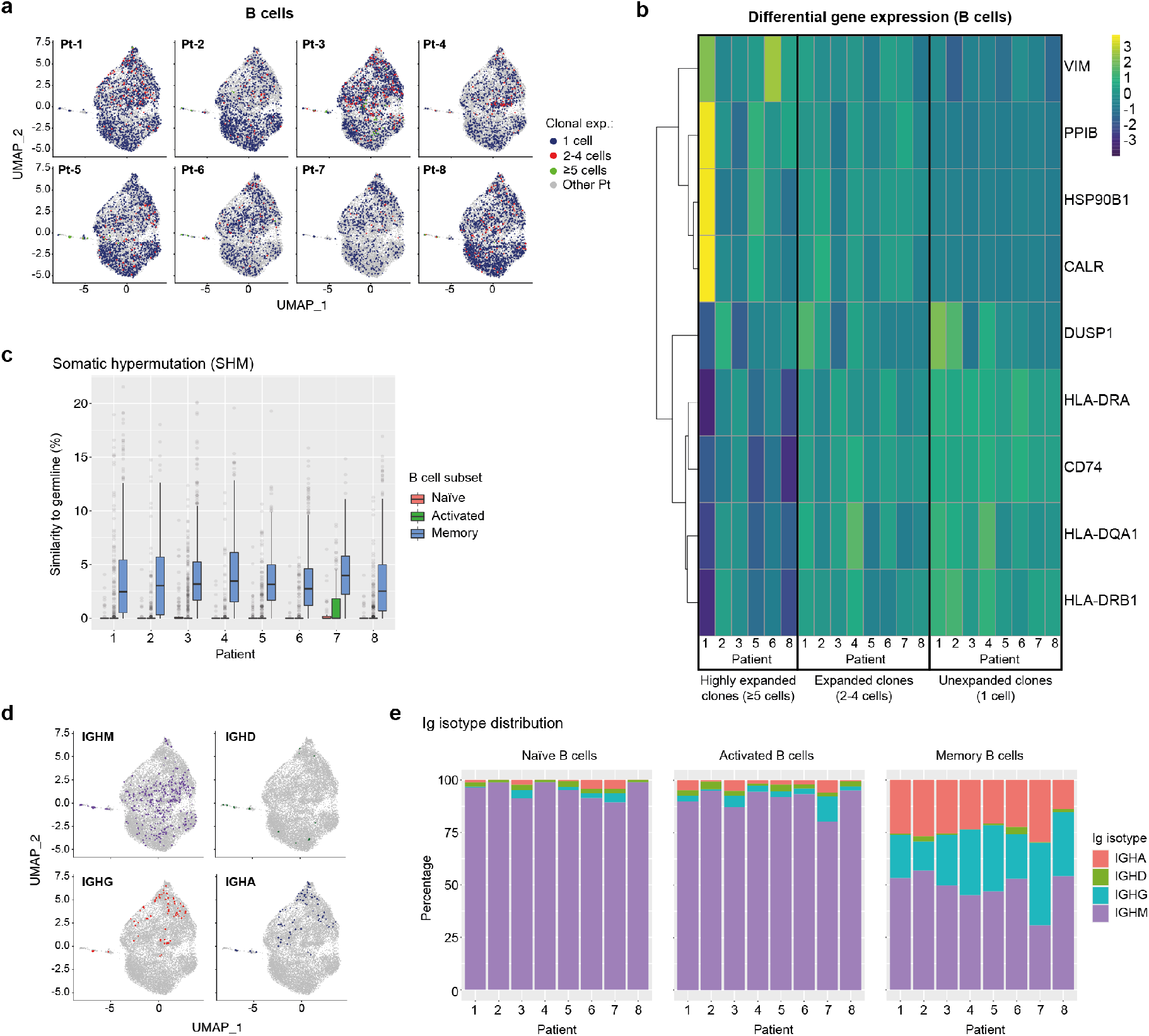
Single-cell transcriptome and BCR profiling reveals elevated class-switching and somatic hypermutation levels in memory B cells. **a**, UMAP plots display B cells from specific patients according to their clonal expansion levels. B cells from other patients in each individual plot are shown in grey. **c**, Heatmap shows differential gene expression (DGE) in unexpanded, expanded and highly expanded B cells. Genes were filtered to include those with detectable expression in at least 50% of cells and that had a minimum 50% fold-change in expression level between groups. **c**, Graph displays the levels of somatic hypermutation (SHM) in the BCRs of naïve, activated and memory B cells across patients. SHM levels are based on the percentage similarity between BCR heavy chain V-gene and its corresponding germline. Data are displayed as median ± IQR. **d**, Graph shows the distribution of B cells expressing specific Ig isotypes relative to their location in transcriptome UMAP plots. B cells from all patients are shown. **e**, Bar graphs show Ig isotype distribution of BCRs found in naïve, activated and memory B cells across patients. CALR = Calreticulin; CD74 = HLA class II Histocompatibility Antigen Gamma Chain; DUSP1 = Dual Specificity Phosphatase 1; HLA-D = Major Histocompatibility Complex, Class II; HSP90B1 = Heat Shock Protein 90 Beta Family Member 1; PPIB = Peptidylprolyl Isomerase B; VIM = Vimentin.

We next analysed single-cell BCR sequencing data to assess SHM levels in different B cell subsets. BCRs from memory B cells showed the highest levels of SHM across patients, while BCRs extracted from naïve and activated B cells displayed similarly low median SHM values. Notably, however, activated B cells expressed a larger number of high-SHM outliers than naïve B cells, suggesting ongoing affinity maturation in this subset (**Fig. 5c**). Mapping of Ig isotype information onto the B cell transcriptional UMAP space, revealed an even distribution of IgM expression across B cell subsets, rare occurrence of IgD-expressing B cells and, most notably, confinement of class-switched IgG- and IgA-expressing B cells to the memory and MZ B cell regions (**Fig. 5d**). Finally, assessment of Ig isotype distribution across patients and B cell subsets revealed that as expected the vast majority of naïve B cells expressed the IgM isotype (**Fig. 5e**), with minimal levels of class-switching observed in activated B cells but prominent class-switching to IgG and IgA isotypes in the memory B cell compartment across all patients (**Fig. 5e**).

Together our results indicate ongoing transition of clonally-expanded B cells into antibody-producing plasma cells, as well as high levels of SHM and Ig isotype class-switching in the memory B cells of convalescent COVID-19 patients.

### Computational prediction of shared specificity identifies candidate SARS-CoV-2-specific TCRs

Motivated by the high levels of CD8+ T cell clonal expansion and activation observed in convalescent COVID-19 patients, we further analyzed single-cell TCR repertoires for potential SARS-CoV-2 specificity. To this end, we applied GLIPH2, an algorithm developed by M. Davis and colleagues that clusters TCRs with a high probability of recognizing the same epitope into specificity groups (based on conserved motifs and similarity levels in CDR3β)^52^. In addition, the provision of HLA typing data enables the prediction of HLA restriction in specific TCR clusters. Analysis of 23,010 paired TCRα and TCRβ sequences derived from the CD8+ T cells of eight patients led to the identification of a total of 552 specificity groups with attributed HLA restriction (seven alleles). We observed distinct proportions of shared specificity groups between pairs of patients, with Pt-1:Pt-8, Pt-3:Pt-7 and Pt-1:Pt-7 showing the highest overlap (**Fig. 6a**). Furthermore, the vast majority of clusters were attributed with HLA-A*01:01, HLA-A*03:01, HLA-B*13:02 or HLA-C*03:04 restriction, and HLA-A*02:01, A*24:02 and C*04:01 were attributed to less than 15% of TCR clusters (**Fig. 6b**). While some of these clusters may be defined by SARS-CoV-2 specificity, it is difficult to exclude reactivity to common human viruses (e.g., CMV, EBV). To further investigate potential for SARS-CoV-2 specificity, we analyzed the sequences of known HLA-A*02:01-restricted SARS-CoV-2-specific, CMV-specific and EBV-specific TCRs alongside those derived from patients expressing the HLA-A*02:01 allele (i.e., Pt-2, Pt-4 and Pt-8) (**Supplementary Tables 3 and 4**). GLIPH2 identified 35 unique patient TCR sequences that clustered together with known SARS-CoV-2-specific TCRs, with the great majority originating from Pt-8 (**Fig. 6c and Table 1**). Thus, such TCRs represent candidates for mediating CD8+ T cell immunity against SARS-CoV-2 infection.

**Figure 6.**
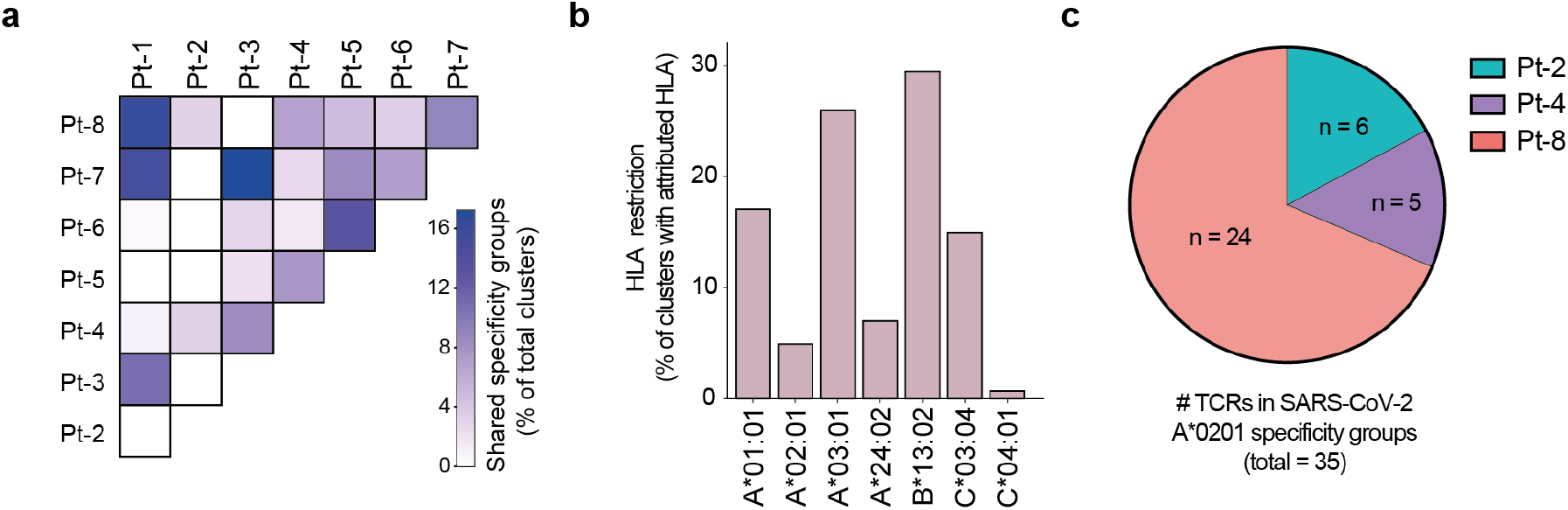
GLIPH2 analysis of single-cell paired TCR repertoires reveals candidate SARS-CoV-2-specific TCRs. **a**, Heatmap shows the proportion of TCR specificity groups containing sequences from specific pairs of patients, as determined by GLIPH2 analysis (total TCR clusters = 552). **b**, Bar plot displays the proportions of predicted HLA class I alleles in HLA-attributed TCR specificity groups (total TCR clusters = 552). **c**, Graph displays the proportions and numbers of candidate SARS-CoV-2-specific TCRs derived from HLA-A*0201-positive patients, as determined by GLIPH2 clustering with known SARS-CoV-2-specific TCR sequences.

### Limitations of the study

Our sample size of eight patients (four per age group) is small and reduces the number of conclusions that we can confidently make from the observed data. Nevertheless, single-cell data offers a deeper characterization of each patient than normal bulk transcriptome and repertoire studies. Furthermore, while the time between symptom onset and sample collection is highly uniform across patients, the time between symptom resolution and sample collection is significantly shorter in the old patient group due to prolonged symptom duration. This is an important variable that should be considered when interpreting the findings presented here.

## Discussion

Here we apply scSeq for in-depth immune repertoire and transcriptomic analysis of T cells and B cells derived from non-severe COVID-19 patients at one month of convalescence. Our analyses of transcriptomic data defined eleven T cell and B cell subsets, of which the effector CD8+ T cell subset (GZMB, NKG7) showed the highest levels of expansion in specific patients, both in terms of proportion and clonality. These findings are in agreement with recent scSeq studies of convalescent COVID-19 patients ^28,53^. For example, effector tissue-resident CD8+ T cells from bronchoalveolar lavage fluid were found to be highly clonally expanded in convalescent COVID-19 patients that experienced moderate but not severe infection^53^. In addition, scSeq of PBMCs revealed increases in cytotoxic effector CD4+ and CD8+ T cell subsets in non-severe convalescent COVID-19 patients only^28^. In our study, we observed high levels of clonal expansion of the effector CD8+ T cell subset but less evident expansion of CD4+ T cell subsets. Importantly, however, differential gene expression analysis revealed that both highly clonally expanded CD8+ and CD4+ T cells had elevated markers of cytotoxicity (CD8: PRF1, GZMH, GNLY; CD4: GZMA).

A recent study of functional T cell responses against SARS-CoV-2 reported significantly higher CD8+ T cell responses directed at spike, M/NP and ORF/Env epitopes in convalescent COVID-19 patients experiencing moderate symptoms compared to those recovering from severe infection^54^. Further evidence supporting a potential role of CD8+ T cells in rapid viral clearance comes from the occurrence of SARS-CoV-2-specific T cells in asymptomatic seronegative family members of COVID-19 patients ^55^, as well as in samples from asymptomatic seronegative control subjects obtained during the COVID-19 pandemic but not prior to it ^55,56^. In line with these findings, our analysis of clonal expansion in lymphocyte subsets suggest a key role of CD8+ effector T cells in the clearance and protection against SARS-CoV-2 in patients with moderate disease. Notably, the fact that younger patients in our cohort had significantly higher CD8-to-CD4 T cell ratios might have contributed to reduced symptom duration relative to older patients. In this context, we predict that methods for the identification of CD8-derived SARS-CoV-2-specific TCRs including functional assays^54,55^ and the application of motif clustering^56^ or machine learning^57^ to TCR repertoire data, as well as methods for the identification of their corresponding epitopes^58,59^ will become increasingly important tools for monitoring SARS-CoV-2 immunity.

In contrast to T cells, we found modest levels of B cell clonal expansion that was not exclusively restricted to a particular subset but spanned activated, memory and MZ B cells. These low levels of B cell clonal expansion are in contrast to the high IgG and IgA serum titers found in all patients of our cohort, particularly those in the older patient group. This suggests that the B cell compartment may experience clonal contraction at one month of convalescence, with the disconnect from serum titers of IgG likely explained by the long half-lives of secreted IgG (~3 weeks)^60^. It should also be noted that antibody-producing plasma cells, which were not included in our analysis, may display clonal expansion levels that are more congruent with the observed levels of SARS-CoV-2-specific antibodies in serum. Although rare, highly expanded B cell clones showed elevated markers of plasma cell transition and activation, thus indicating possible ongoing differentiation into plasma cells, albeit at low levels, at one month after symptom onset.

Analysis of BCR repertoires and transcriptomes revealed that the highest levels of SHM and Ig class-switching occurred in the memory B cell subset. This finding is consistent with the generation of germinal center-derived memory B cells following antigen encounter^61,62^. Similarly, SHM and Ig class-switching levels were directly correlated with B cell clonal expansion. Remarkably, we found that highly expanded B cell clones from the young patient group had significantly higher SHM levels (median = 6.7% divergence from germline) than those derived from the old patient group (median = 2.5% divergence from germline). This occurred despite significantly higher serum titers of SARS-CoV-2-specific antibodies in the old patient group, and suggests that more effective affinity maturation may occur in younger patients. Initial studies characterizing SARS-CoV-2-specific antibodies reported low levels of SHM, with median divergence from germline ranging from 0.7-2% in convalescent patients of varied symptom severity analyzed at 20-40 days following symptom onset^63–65^. Notably, a recent report has described ongoing affinity maturation of SARS-CoV-2-specific antibodies at six months following symptom onset in non-severe patients, with SHM levels rising to 3% divergence from germline^2^. Interestingly, the cited study provides evidence of viable SARS-CoV-2 antigen in the gut of such patients, which has been proposed as a source of antigen for ongoing affinity maturation linked to elevated IgA serum titers. It is thus unclear whether increased SHM levels found in the young patients of our cohort are the result of a better capacity for affinity maturation upon initial antigen encounter or of ongoing affinity maturation resulting from longer exposure to antigen following symptom resolution. The clear dominance of IgA class-switched BCRs expressed by highly expanded B cells in 3 out of 4 patients in the young group appears to support the latter.

In conclusion, our in-depth characterization integrating single-cell immune repertoire and transcriptome profiling of T and B cells represents a valuable resource to better understand the adaptive immune response and age-related differences in convalescent COVID-19 patients with moderate disease. Furthermore, it serves as an important point of reference for future single-cell characterization of lymphocytes at later time points of convalescence, or of lymphocytes isolated from patients experiencing long-term COVID-19 sequelae and can help in defining markers for clinical monitoring of disease progression.

## Methods

### Patient samples

Patients were participants of the SERO-BL-COVID-19 study sponsored by the Department of Health, Canton Basel-Landschaft, Switzerland. All analyzed patients tested positive for SARS-CoV-2 after RT-PCR of naso- and oropharyngeal swab samples and experienced a resolution of COVID-19 symptoms without requiring hospitalization. Whole blood was collected 25 to 39 days following a positive RT-PCR test and subjected to density gradient centrifugation using the Ficoll Paque Plus reagent (GE Healthcare, #17-1440-02). After separation, the upper plasma layer was collected for ELISA detection of IgG and IgA SARS-CoV-2-specific antibodies (Euroimmun Medizinische Labordiagnostika, #EI2668-9601G, #EI2606-9601A). Peripheral blood mononuclear cells (PBMC) were collected from the interphase, resuspended in freezing medium (RPMI 1640, 10%(v/v) FBS, 10%(v/v) dimethyl sulfoxide) and cryopreserved in liquid nitrogen. Point-of-care lateral flow immunoassays assessing the presence of IgG and IgM SARS-CoV-2-specific antibodies (Qingdao Hightop Biotech, #H100) were performed at the time of blood collection.

### Immunomagnetic isolation of B cells and T cells

PBMC samples were thawed, washed in complete media (RPMI 1640, 10%(v/v) FBS) and pelleted by centrifugation. Cells were resuspended in 0.5 mL complete media, counted and treated with 10 U ml^−1^ DNAse I (Stemcell Technologies, #) for 15 min at RT in order to prevent cell clumping. After DNase I digestion, cells were washed twice in complete media, pelleted by centrifugation and resuspended in 0.5 mL flow cytometry buffer (PBS, 2%(v/v) FBS, 2 mM EDTA). The cell suspension was filtered through a 40 μM cell strainer prior to immunomagnetic isolation. As a first step, plasma cells were isolated using the EasySep Human CD138 Positive Selection Kit II (Stemcell Technologies, #17877) for analysis in a companion study (manuscript in preparation). The negative fraction of the above selections was divided into two aliquots that were subjected to negative immunomagnetic isolation of either B cells (EasySep Human Pan-B cell Enrichment Kit, Stemcell Technologies, #19554) or T cells (EasySep Human T cell Isolation Kit, Stemcell Technologies, #17951). After isolation, B cells and T cells were pelleted by centrifugation, resuspended in PBS, 0.4%(v/v) BSA, filtered through a 40 μM cell strainer and counted. T cells and B cells originating from the same patient were pooled in equal numbers and the final suspension was counted and assessed for viability using a fluorescent cell counter (Cellometer Spectrum, Nexcelom). Whenever possible, cells were adjusted to a concentration of 1×10^6^ live cells/mL in PBS, 0.04%(v/v) BSA before proceeding with droplet generation.

### Single cell droplet generation and preparation of sequencing libraries

Encapsulation of lymphocytes and DNA-barcoded gel beads was performed using the Chromium controller (10x Genomics, PN-110203). Briefly, 1.4×10^4^ to 1.7×10^4^ cells (in reverse transcription mix) were loaded per channel for a targeted recovery of 8×10^3^ to 1×10^4^ cells per sample. Reverse transcription and preparation of single-cell transcriptome, BCR and TCR libraries was performed according to the manufacturer’s instructions (CG000086 manual, RevM, 10x Genomics) and using the following kits: Chromium Single Cell 5’ Library & Gel Bead Kit (PN-1000006), Chromium Single Cell 5’ Library Construction Kit (PN-1000020), Chromium Single Cell V(D)J Enrichment Kit, Human T Cell (PN-1000005), Chromium Single Cell V(D)J Enrichment Kit, Human B Cell (PN-1000016), Chromium Single Cell A Chip Kit (PN-1000009), Chromium i7 Multiplex Kit (PN-120262).

### Deep sequencing

The quality and concentrations of transcriptome (i.e., cDNA), TCR and BCR libraries were determined using a fragment analyzer (Agilent Bioanalyzer) at specific steps of library preparation, as recommended in the 10x Genomics scSeq protocol (CG000086 manual, RevM). Following multiplexing (Chromium i7 Multiplex Kit, #PN-120262, 10x Genomics), transcriptome libraries were treated with free adapter blocking (FAB) reagent to prevent index switching (#20024144, Illlumina). Paired-end sequencing of multiplexed transcriptome libraries was performed using a NovaSeq 6000 sequencer (Illumina) and SP100-cycle kit (#20027464, Illumina). TCR and BCR libraries were multiplexed, FAB-treated and paired-end-sequenced using a second SP100-cycle kit in a separate run.

### HLA class I typing

HLA class I transcripts were amplified and deep-sequenced from two overlapping RT-PCR reactions flanking exons 2 (PCR 1) or 3 (PCR 2) using barcoded primers designed to target conserved regions (**Supplementary Table 5**)^66^. Total RNA was extracted from patient PBMCs by resuspension in TRIzol reagent (Invitrogen, # 15596018), and column-purified using the PureLink RNA Mini kit (Invitrogen, #12183025). For reverse transcription, 100 pmol of oligo dT, 10 nmol of each dNTP, 40 ng RNA and sufficient nuclease-free water for a final 14 μl volume were mixed, incubated at 65°C for 5 min and chilled on ice for 5 min. This was followed by addition of 4 μL of 5X RT buffer, 40 units of RiboLock RNAse inhibitor (Thermo Fisher, #EO0381) and 200 units of Maxima H-minus reverse transcriptase (Thermo Fisher, #EP0751) and mixing. Reverse transcription was performed at 50°C for 30 min, followed by inactivation at 85°C for 5 min. 5 μl of the resulting cDNA-containing reverse transcription mixes were then used as templates for 25 μL PCR reactions using the KAPA HiFi PCR kit with GC buffer (Roche Diagnostics, #07958846001) and the following thermal cycling conditions: 95°C for 3 min; 35 cycles of 98°C for 20 s, 61°C for 15 s, 72°C for 15 s; and final extension at 72°C for 30 s. HLA amplicons were purified by gel-extraction (QIAquick gel extraction kit, Qiagen #28704) and submitted for Illumina paired-end deep-sequencing (Amplicon-EZ, Genewiz). Unique sequences originating from specific patients were identified from their respective DNA barcodes and aligned using the ClustalOmega tool to cluster sequences arising from the same allele. Sequences with the highest amount of reads in each cluster were used as input for the basic local alignment search tool (BLAST; Nucleotide collection, *Homo sapiens*). Sequences returning matching or highly similar alleles across PCR 1 and PCR 2 in each patient were then assembled and queried against the IMGT/HLA database for final validation.

### Transcriptome scSeq alignment and quality control (QC)

Reads from transcriptome scSeq (FASTQ format) were aligned to the GRCh38 reference human genome and output as filtered gene expression matrices using the 10x Genomics Cell Ranger software (version 3.1.0). Subsequent data QC and analysis was performed using R (version 3.6.2) and the Seurat package (version 3.1.5). QC steps consisted of the exclusion of TCR and BCR genes (prevention of clonotype influence on subsequent clustering), the exclusion of cells with lower than 150 or greater than 3500 genes (low quality cells), and the exclusion of cells in which more than 20% of UMIs were associated with mitochondrial genes (reduction of freeze-thaw metabolic effects)^67^.

### Dataset normalisation and integration of multiple datasets

Patient datasets were merged into a Seurat object list using the *merge* and *SplitObject* function. Each patient dataset was then separately normalised using *SCTransform*. Variable integration features (3,000) were calculated using the *SelectIntegrationFeatures* function from the R package Seurat^68^ and setting them as variable features after merging the normalised patient datasets. Principal component analysis for dimensionality reduction was performed using the *RunPCA* function with up to 50 principal components. Potential batch effects between patient samples were addressed with the Harmony R package (version 1.0) using the *RunHarmony* function ^69^. Finally, unsupervised clustering was performed using the *FindNeighbours* and *FindClusters* functions. Non-linear dimensionality reduction using the *RunUMAP* function was performed using the first 50 principal components to generate the final UMAP visualization of cell clusters.

### Dataset subsetting of CD8+ T cells, CD4+ T cells and B cells

Initial T and B cell separation was performed by mapping of TCR and BCR (VDJ) cell-specific barcodes onto the scSeq transcriptome dataset. Double attribution of TCR and BCR to the same cell (i.e., barcode) was used to identify and exclude doublets. Separation of CD8+ and CD4+ T cells was performed using the *WhichCells* function, from the R package Seurat, based on the singular expression of CD8A and CD4, respectively. Additional filtering of B cells was done by discarding all B cells that showed expression of CD3E or SDC1 as well as excluding B cells whose cellular barcodes occured in the Plasma cell BCR (VDJ) cell barcodes (data not shown).

### Cell state annotation and marker identification

The expression of specific markers in identified clusters was determined using the *FindAllMarkers* function using the Wilcoxon Rank Sum test. Cluster-specific markers were thresholded by having a log2(fold-change) greater than 0.25 between cells in the respective cluster and remaining cells; with marker expression occurring in at least 25% of cells in the cluster. Clusters were then attributed with specific cell states based on the expression of canonical markers

### Differential gene expression analysis

Differentially expressed genes between two groups of cells were identified using the *FindMarkers* function. Genes were thresholded by being expressed in more than 50% of the cells and by having a log2(fold-change) greater than 0.5 between cells of the different groups using the Wilcoxon Rank Sum test.

### Paired TCR and BCR (VDJ) single-cell sequencing alignment and QC

TCR and BCR reads in FASTQ format were aligned with the VDJ-GRCh38-alts-ensembl reference using the 10x Genomics Cell Ranger VDJ software (version 3.1.0). This generated single-cell VDJ sequences and annotations such as gene usage, clonotype frequency and cell-specific barcode information. As a QC step, only cells with one productive alpha and one productive beta chain (T cells) or with one productive heavy and one productive light chain (B cells)were retained for downstream analysis.

### Paired TCR and BCR (VDJ) analysis

Clonotype definition was adjusted to count all sequences as clonal if they met the following criteria: (1) Same V and J gene usage in both chains, (2) Same CDR3 length in both chains and (3) 80% amino acid sequence similarity in the CDR3 region of the TCRβ (T cells) or BCR heavy chain (B cells). Shared cellular barcode information between TCR/BCR (VDJ) scSeq and transcriptome scSeq data was used to project TCR and BCR clonotypes onto the UMAP plots (colour-coded by clonal expansion level).

### Somatic hypermutation analysis

SHM levels in individual BCR clonotypes were determined using the change-o toolkit from the Immcantation portal as a wrapper to run IgBlast on the Cell Ranger VDJ output. The IgBlast output enabled assessment of germline similarity of single-cell BCR (VDJ) sequences. Germline identity was used as a proxy for somatic hypermutation levels and was calculated from alignments of BCR clonotypes with their corresponding VH and VL germline sequences.

### TCR specificity group identification using GLIPH2

GLIPH2 clusters TCRs into specificity groups predicted to share the same antigen specificity based on sequence similarity^52^. We used this algorithm to cluster TCRs from HLA-A*0201 patients (Pt-2, Pt-4 and Pt-8) together with known SARS-CoV-2 binders as well as CMV and EBV binders, which were also from HLA-A*0201 background (obtained from VDJdb database). Specificity groups that were reported by GLIPH2, were filtered for groups that were significant according to the Fisher’s Exact test (significance level < 5%) and contained at least one patient TCR and one TCR of known specificity (i.e., SARS-CoV-2, EBV or CMV). Specificity groups were identified with either global (0-1 amino acid differences in CDR3β) or local similarities (CDR3β share a common motif that is rare in the reference dataset). CD4 expressing clonotypes were filtered out.

### Pseudotime analysis

Pseudotime and trajectory inference was applied to scSeq transcriptome data using the *slingshot* function with default parameters from the Slingshot package in R^70^. The naive cluster from each CD8+ T cell, CD4+ T cell and B cell subgroup was set as the starting point for the minimum spanning tree. The previously generated UMAP clustering was set as the cellular embedding on which Slingshot performed trajectory inference computation.

### Code availability

The data analysis pipeline followed the standard procedures as outlined in Cell Ranger and Seurat documentations. Custom scripts and functions for easier downstream analysis and visualisation purposes are available upon request.

### List of utilized R packages

Biobase (2.46.0), BiocGenerics (0.32.0), BiocParallel (1.20.1), <u>Cell Ranger (3.1.0)</u>, Change-O (1.0.0), circlize (0.4.10), data.table (1.12.8), DelayedArray (0.12.3), dplyr (0.8.5), GenomeInfoDb (1.22.1), GenomicRanges (1.38.0), ggplot2 (3.3.2.9000), harmony (1.0), pheatmap (1.0.12), princurve (2.1.5), RColorBrewer (1.1-2), matrixStats (0.56.0), sctransform (0.2.1), <u>Seurat (3.1.5)</u>, slingshot (1.4.0), stringdist (0.9.5.5), stringr (1.4.0), tibble (3.0.3), tidyr (1.1.0), tidyverse (1.3.0).

## Supplementary Information

Supplementary Figs. 1-7, Supplementary Tables 1-5.

## Acknowledgements

We thank the Genomics Facility Basel (D-BSSE, ETH Zurich), in particular Ms. Ina Nissen and Dr. Christian Beisel, for their excellent support with the scSeq protocol and for performing deep sequencing runs.

## Author contributions

F.B., R.V.-L., A.Y. and S.T.R. designed the study; F.B., R.V.-L., R.A.E., D.M.M., B.W., E.K., R.B.D.R. performed experiments; F.B., R.V.-L., A.Y., C.R.W. analyzed data; F.B., R.V.-L., A.Y. and S.T.R. wrote the manuscript with input from all authors.

## Data and Materials Availability

The raw FASTQ files from deep sequencing that support the findings of this study will be deposited under (DOI link from zenodo). Additional data that support the findings of this study are available from the corresponding author upon reasonable request.

