## Supplementary Material for "A single-cell atlas of lymphocyte adaptive immune repertoires and transcriptomes reveals age-related differences in convalescent COVID-19 patients"

### Supplementary Information

Supplementary Figures 1-7

Supplementary Tables 1-5

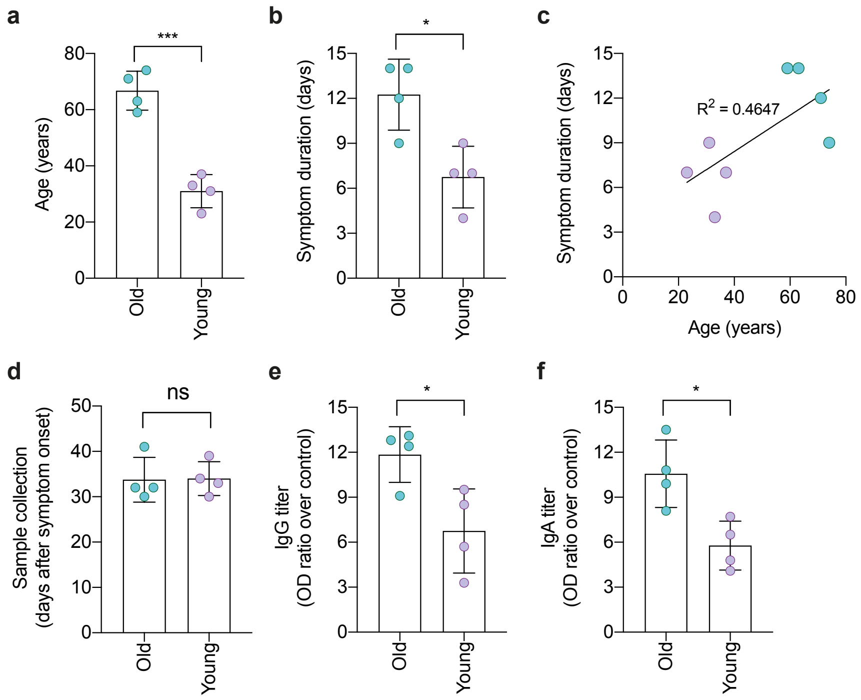


**Supplementary Figure 1. Clinical and serological characteristics of convalescent COVID-19 patients**. **a-b**, Age and COVID-19 symptom duration in patients from the old and young groups. **c**, Linear regression analysis of age and symptom duration in convalescent COVID-19 patients. **d-f**, Serological analysis of convalescent COVID-19 patients displaying time of blood collection (**d**), SARS-CoV-2-specific IgG antibody titers (**e**) and SARS-CoV-2-specific IgA antibody titers (**f**), as determined by ELISA. Data are displayed as mean ± SD. Asterisks indicate significant differences between groups. * $P < 0.05$, ** $P < 0.01$, *** $P < 0.001$, **** $P < 0.0001$, ns = not significant.

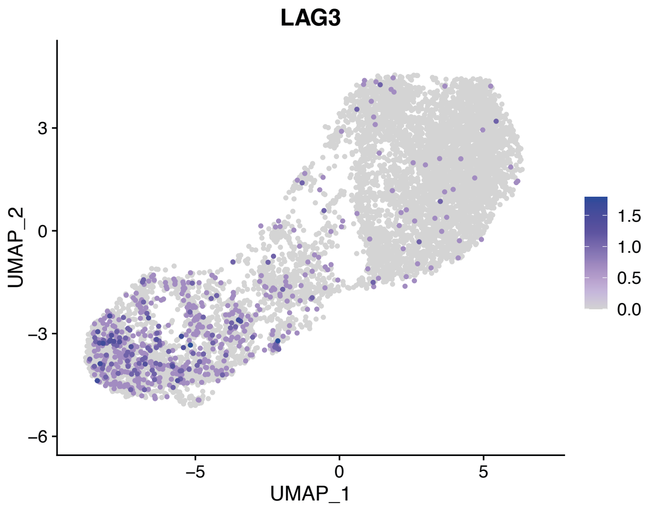


**Supplementary Figure 2. Exhausted CD8+ T cells predominantly localize to the effector CD8+ T cell region**. UMAP plot shows expression of the LAG-3 exhaustion marker by individual CD8+ T cells from all patients.

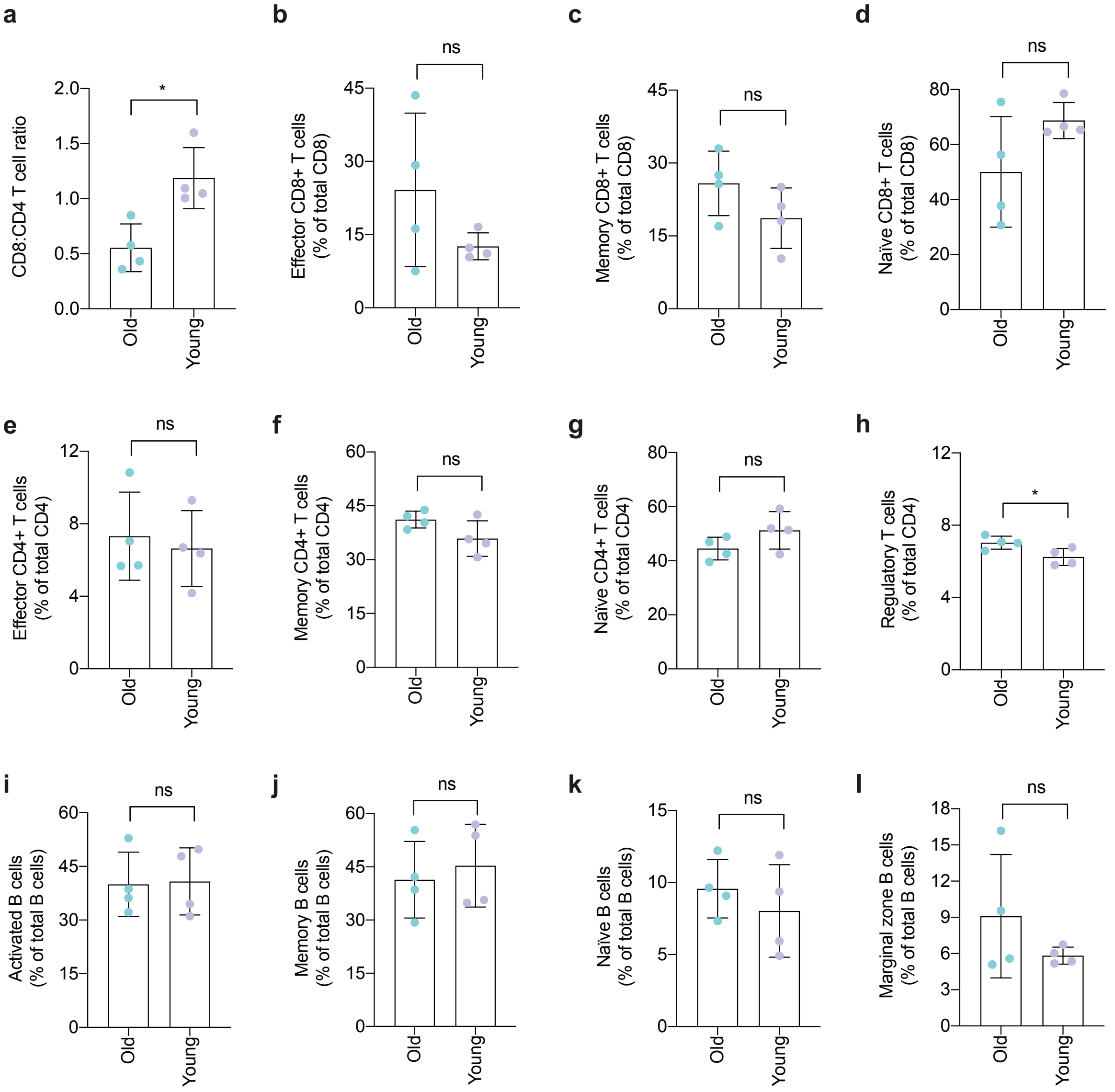


**Supplementary Figure 3. Proportions of lymphocyte subsets across studied age groups.** Bar graphs compare the proportions of T cell and B cell subsets in the old and young groups. Data are displayed as mean ± SD. Asterisks indicate significant differences between groups. * P < 0.05, ** P < 0.01, *** P < 0.001, **** P < 0.0001, ns = not significant.

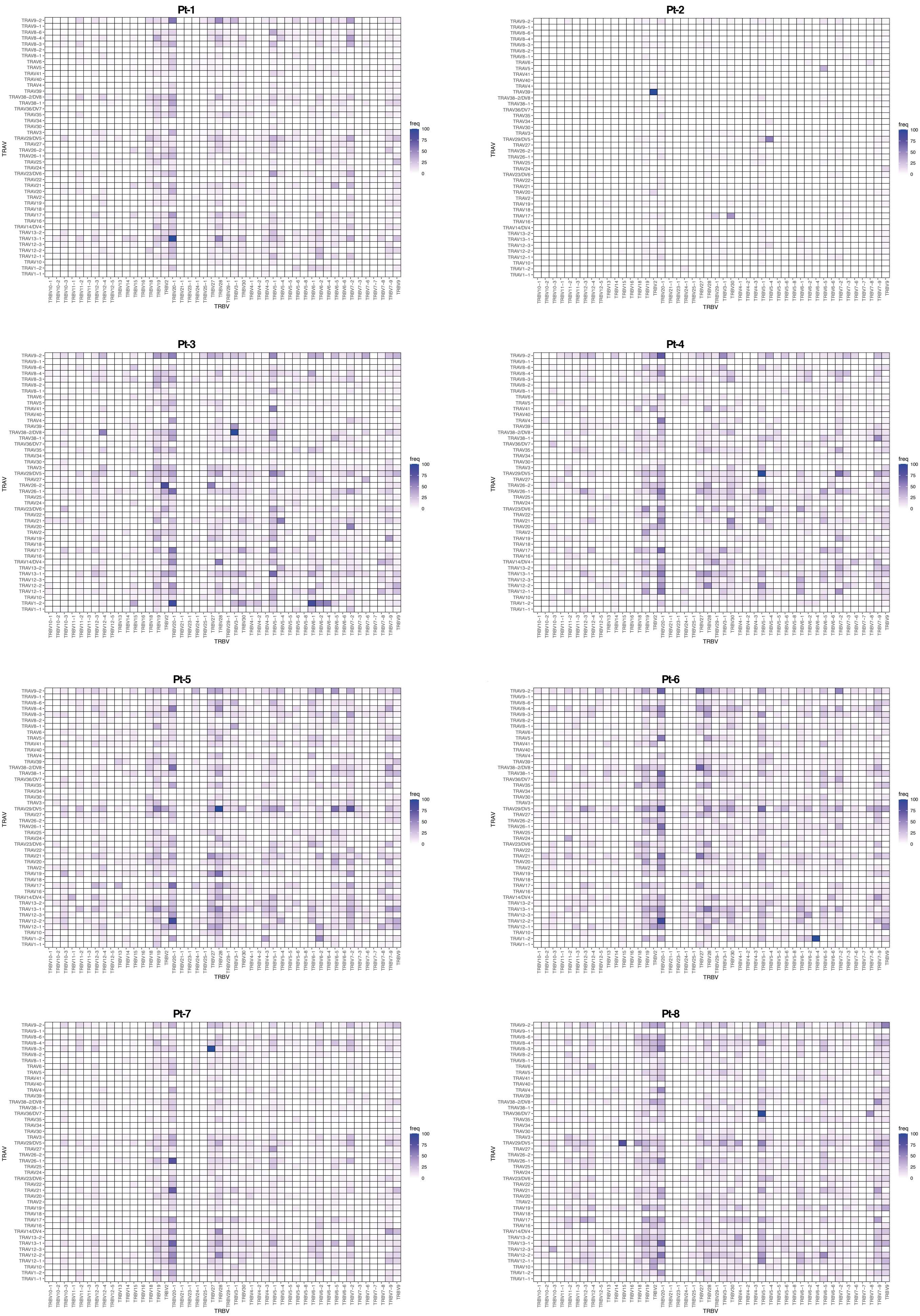
Pt-1
Pt-2
Pt-3
Pt-4
Pt-5
Pt-6
Pt-7
Pt-8
TRAV
TRBV
freq
100
75
50
25
0

**Supplementary Figure 4**. **TCR V-gene pairings in convalescent COVID-19 patients.** Heatmaps display the relative frequencies of specific TCR V-gene pairings in the T cells of individual patients. Data is normalized to the most frequent pairing found in each patient.

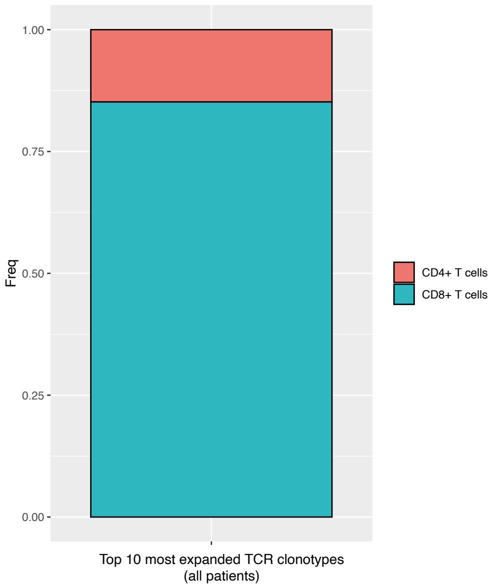


**Supplementary Figure 5**. **T cell clonal expansion in convalescent COVID-19 patients is dominated by CD8+ subsets**. Bar graph shows the proportions of CD8+ T cells versus CD4+ T cells within the top ten most expanded TCR clonotypes from each patient.

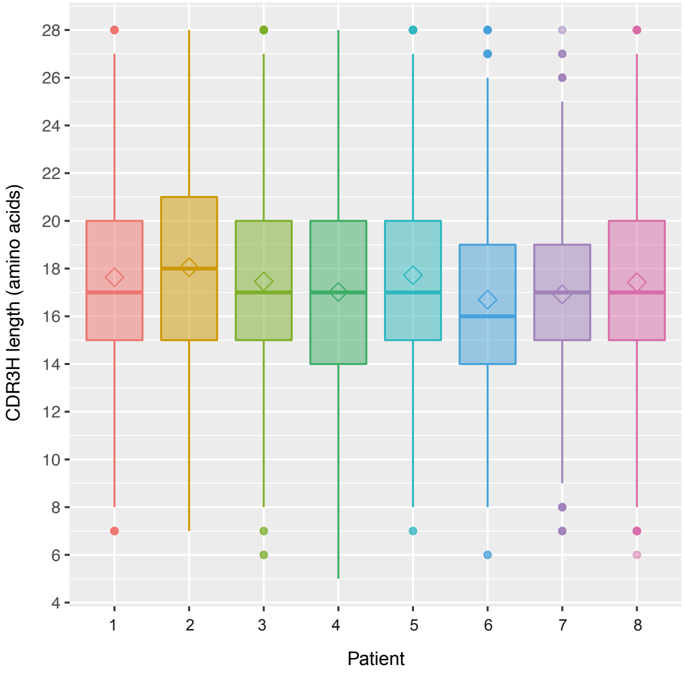


**Supplementary Figure 6**. **BCR heavy chain CDR3 length distribution in convalescent COVID-19 patients.**

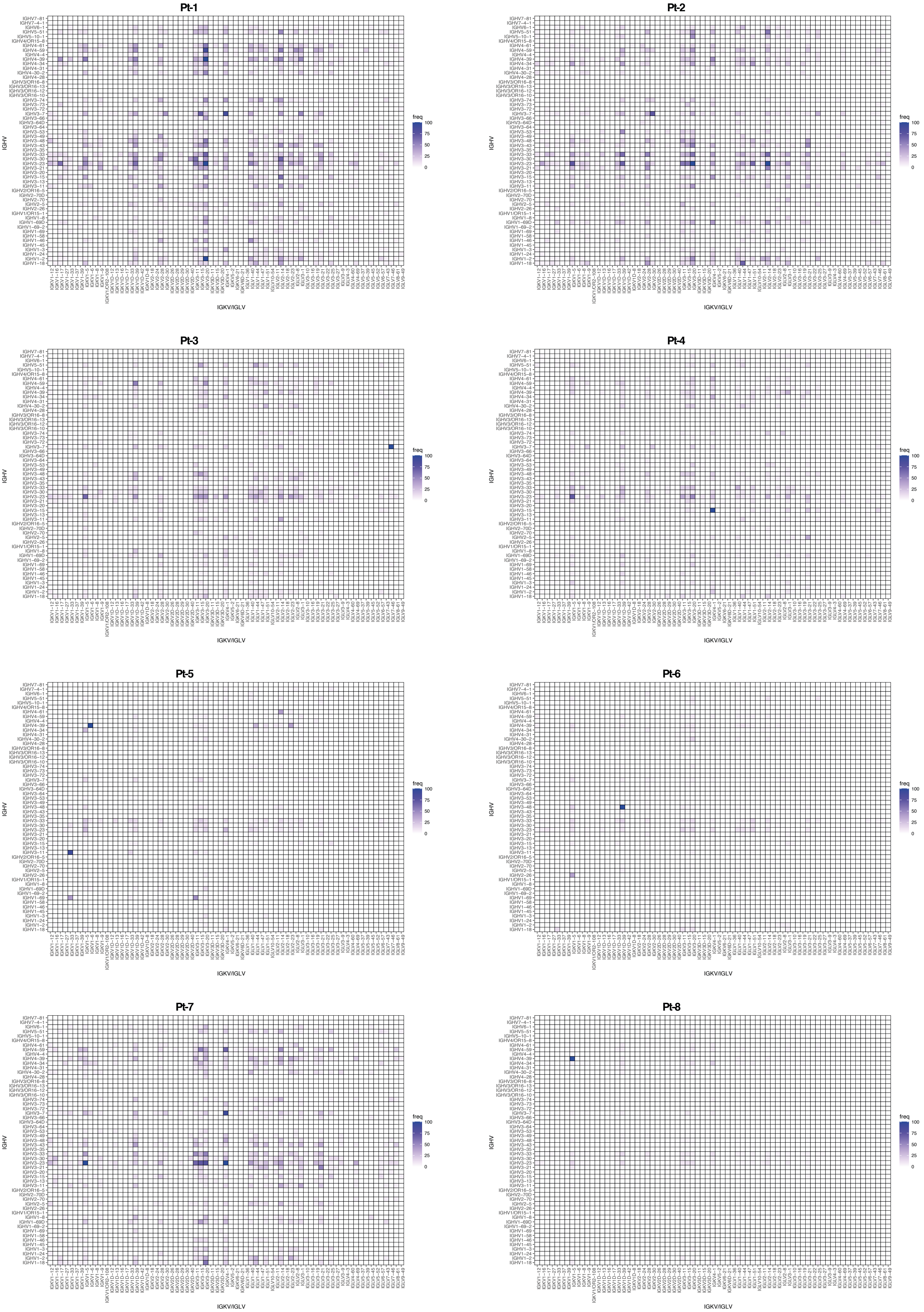
Pt-1
Pt-2
Pt-3
Pt-4
Pt-5
Pt-6
Pt-7
Pt-8
IGHV
IGKV/IGLV
freq
100
75
50
25
0

**Supplementary Figure 7. BCR V-gene pairings in convalescent COVID-19 patients.** Heatmaps display the relative frequencies of specific BCR V-gene pairings in the B cells of individual patients. Data is normalized to the most frequent pairing found in each patient.

**Supplementary Table 1.** Characteristics of convalescent COVID-19 patients analyzed in this study

| Patient ID | Age | Sex | BMI ($kg/m^2$) | RT-PCR test | Days ill (before PCR / after PCR / total) | Restrictions | Sample collection (days after symptom onset) | Rapid test (LFA) | ELISA test | IgA titer (OD ratio) | IgG titer (OD ratio) |
|---|---|---|---|---|---|---|---|---|---|---|---|
| Pt-1 | 59 | F | 33.9 | Positive | 7 / 7 / 14 | Bedridden | 32 | IgM+ / IgG+ | IgA+ / IgG+ | 10.8 | 13.1 |
| Pt-2 | 63 | M | 24.4 | Positive | 7 / 7 / 14 | Bedridden | 41 | IgM+ / IgG+ | IgA+ / IgG+ | 13.5 | 12.8 |
| Pt-3 | 74 | F | 23.4 | Positive | 2 / 7 / 9 | Help needed | 32 | IgM+ / IgG+ | IgA+ / IgG+ | 8.1 | 9.1 |
| Pt-4) | 71 | F | 28.9 | Positive | 5 / 7 / 12 | Bedridden | 30 | IgM+ / IgG+ | IgA+ / IgG+ | 9.9 | 12.4 |
| Pt-5 | 37 | M | 28.7 | Positive | 2 / 5 / 7 | No restrictions | 33 | IgM- / IgG+ | IgA+ / IgG+ | 7.7 | 3.3 |
| Pt-6 | 31 | M | 23.2 | Positive | 2 / 7 / 9 | No restrictions | 34 | IgM- / IgG+ | IgA+ / IgG+ | 6.5 | 8.5 |
| Pt-7 | 23 | F | 22 | Positive | 0 / 7 / 7 | No restrictions | 39 | IgM+ / IgG+ | IgA+ / IgG+ | 4.8 | 5.7 |
| Pt-8 | 33 | M | 26 | Positive | 2 / 2 / 4 | No restrictions | 30 | IgM+ / IgG+ | IgA+ / IgG+ | 4.1 | 9.5 |

**Supplementary Table 2.** HLA class I alleles of studied convalescent COVID-19 patients

| Patient | HLA-A | HLA-B | HLA-Cw |
|---|---|---|---|
| Pt-1 | A*03:01, A*30:01 | B*13:02, B*40:02 | C*02:02, C*06:02 |
| Pt-2 | A*02:01, A*26:01 | B*38:01, B*44:02 | C*05:01, C*12:03 |
| Pt-3 | A*01:01, A*24:02 | B*07:02, B*15:29 | C*07:02, C*07:04 |
| Pt-4 | A*02:01, A*02:01 | B*15:01, B*35:01 | C*03:04, C*04:01 |
| Pt-5 | A*11:01, A*11:01 | B*18:03, B*40:01 | C*03:04, C*07:01 |
| Pt-6 | A*01:01, A*68:01 | B*07:02, B*35:03 | C*04:01, C*07:02 |
| Pt-7 | A*03:01, A*24:02 | B*18:01, B*35:02 | C*04:01, C*07:01 |
| Pt-8 | A*02:01, A*24:02 | B*13:02, B*35:02 | C*04:01, C*06:02 |

**Supplementary Table 3.** GLIPH2 analysis of patient TCRs and known HLA-A*02:01-restricted SARS-CoV-2-specific TCRs

| Cluster | Motif | CDR3b | TRBV | TRBJ | CDR3a | TRAV | TRAJ | Clone | Patient |
|---|---|---|---|---|---|---|---|---|---|
| 1 | QNTG | CASSmaraqNTGELFF | TRBV6-6 | TRBJ2-2 | CAGSIGFGNVLHC | TRAV25 | TRAJ35 | 1059 | Pt-02 |
| 1 | QNTG | CASrdsqNTGELFF | TRBV25-1 | TRBJ2-2 | CAVNSYGGSQGNLIF | TRAV12-2 | TRAJ42 | 20 | Pt-08 |
| 1 | QNTG | CASidqNTGELFF | TRBV5-1 | TRBJ2-2 | NA | NA | NA | NA | NA |
| 1 | QNTG | CASmdqNTGELFF | TRBV2 | TRBJ2-2 | NA | NA | NA | NA | NA |
| 1 | QNTG | CASSYqNTGELFF | TRBV7-8 | TRBJ2-2 | NA | NA | NA | NA | NA |
| 1 | QNTG | CASqdqNTGELFF | TRBV19 | TRBJ2-2 | NA | NA | NA | NA | NA |
| 1 | QNTG | CASSfqNTGELFF | TRBV7-8 | TRBJ2-2 | NA | NA | NA | NA | NA |
| 1 | QNTG | CAvLAeqNTGELFF | TRBV10-2 | TRBJ2-2 | NA | NA | NA | NA | NA |
| 1 | QNTG | CASnnqNTGELFF | TRBV7-8 | TRBJ2-2 | NA | NA | NA | NA | NA |
| 1 | QNTG | CASGtqNTGELFF | TRBV5-1 | TRBJ2-2 | NA | NA | NA | NA | NA |
| 1 | QNTG | CASlsqNTGELFF | TRBV7-8 | TRBJ2-2 | NA | NA | NA | NA | NA |
| 1 | QNTG | CATqvdqNTGELFF | TRBV5-1 | TRBJ2-2 | NA | NA | NA | NA | NA |
| 1 | QNTG | CSVEeqNTGELFF | TRBV29-1 | TRBJ2-2 | NA | NA | NA | NA | NA |
| 1 | QNTG | CARgdqNTGELFF | TRBV7-8 | TRBJ2-2 | NA | NA | NA | NA | NA |
| 1 | QNTG | CASSSqNTGELFF | TRBV7-8 | TRBJ2-2 | NA | NA | NA | NA | NA |
| 1 | QNTG | CASGrqNTGELFF | TRBV5-1 | TRBJ2-2 | NA | NA | NA | NA | NA |
| 1 | QNTG | CAptpdqNTGELFF | TRBV12-3 | TRBJ2-2 | NA | NA | NA | NA | NA |
| 1 | QNTG | CASnsqNTGELFF | TRBV12-3 | TRBJ2-2 | NA | NA | NA | NA | NA |
| 1 | QNTG | CASSVqNTGELFF | TRBV7-8 | TRBJ2-2 | NA | NA | NA | NA | NA |

| | | | | | | | | | |
|---|---|---|---|---|---|---|---|---|---|
| 1 | QNTG | CSARdeaGGQNTGELFF | TRBV20-1 | TRBJ2-2 | NA | NA | NA | NA | NA |
| 1 | QNTG | CSARdqpGGQNTGELFF | TRBV20-1 | TRBJ2-2 | NA | NA | NA | NA | NA |
| 1 | QNTG | CSARdqaaqNTGELFF | TRBV20-1 | TRBJ2-2 | NA | NA | NA | NA | NA |
| 1 | QNTG | CSARdLAaqNTGELFF | TRBV20-1 | TRBJ2-2 | NA | NA | NA | NA | NA |
| 1 | QNTG | CSARdrLAqNTGELFF | TRBV20-1 | TRBJ2-2 | NA | NA | NA | NA | NA |
| 1 | QNTG | CSARsrQGqNTGELFF | TRBV20-1 | TRBJ2-2 | NA | NA | NA | NA | NA |
| 1 | QNTG | CSARgesdqNTGELFF | TRBV20-1 | TRBJ2-2 | NA | NA | NA | NA | NA |
| 1 | QNTG | CASSDttrqNTGELFF | TRBV2 | TRBJ2-2 | NA | NA | NA | NA | NA |
| 1 | QNTG | CASrsrveqNTGELFF | TRBV2 | TRBJ2-2 | NA | NA | NA | NA | NA |
| 1 | QNTG | CSARgGLsqNTGELFF | TRBV20-1 | TRBJ2-2 | NA | NA | NA | NA | NA |
| 1 | QNTG | CSARgGQGqNTGELFF | TRBV20-1 | TRBJ2-2 | NA | NA | NA | NA | NA |
| 1 | QNTG | CSARdGLAqNTGELFF | TRBV20-1 | TRBJ2-2 | NA | NA | NA | NA | NA |
| 1 | QNTG | CASrGLAeqNTGELFF | TRBV2 | TRBJ2-2 | NA | NA | NA | NA | NA |
| 1 | QNTG | CASrdihqqNTGELFF | TRBV2 | TRBJ2-2 | NA | NA | NA | NA | NA |
| 1 | QNTG | CSARdsraqNTGELFF | TRBV20-1 | TRBJ2-2 | NA | NA | NA | NA | NA |
| 1 | QNTG | CSARdDRaqNTGELFF | TRBV20-1 | TRBJ2-2 | NA | NA | NA | NA | NA |
| 1 | QNTG | CSARdaeGQNTGELFF | TRBV20-1 | TRBJ2-2 | NA | NA | NA | NA | NA |
| 3 | DIE | CASSLadierETQYF | TRBV11-2 | TRBJ2-5 | CAVRDQAGTALIF | TRAV2 | TRAJ15 | 1420 | Pt-08 |
| 3 | DIE | CASirdieiNEQFF | TRBV28 | TRBJ2-1 | CVVFNQAGTALIF | TRAV12-1 | TRAJ15 | 2740 | Pt-08 |
| 3 | DIE | CASSLdietYF | TRBV7-9 | TRBJ2-7 | NA | NA | NA | NA | NA |
| 3 | DIE | CASSwdiEAFF | TRBV7-9 | TRBJ1-1 | NA | NA | NA | NA | NA |
| 3 | DIE | CASTrdiEAFF | TRBV7-9 | TRBJ1-1 | NA | NA | NA | NA | NA |

| | | | | | | | | | |
|---|---|---|---|---|---|---|---|---|---|
| 3 | DIE | CASSVdiEAFF | TRBV7-9 | TRBJ1-1 | NA | NA | NA | NA | NA |
| 3 | DIE | CASTqdiEAFF | TRBV7-9 | TRBJ1-1 | NA | NA | NA | NA | NA |
| 3 | DIE | CASStdiEAFF | TRBV7-9 | TRBJ1-1 | NA | NA | NA | NA | NA |
| 3 | DIE | CASSSdiEQFF | TRBV7-9 | TRBJ2-1 | NA | NA | NA | NA | NA |
| 3 | DIE | CASSSdiEAFF | TRBV7-9 | TRBJ1-1 | NA | NA | NA | NA | NA |
| 3 | DIE | CASSDdiEAFF | TRBV7-9 | TRBJ1-1 | NA | NA | NA | NA | NA |
| 3 | DIE | CASSQdiEQYF | TRBV7-9 | TRBJ2-7 | NA | NA | NA | NA | NA |
| 3 | DIE | CASSQdiEAFF | TRBV7-9 | TRBJ1-1 | NA | NA | NA | NA | NA |
| 3 | DIE | CASSPdiEQFF | TRBV7-9 | TRBJ2-1 | NA | NA | NA | NA | NA |
| 3 | DIE | CASSPdiEQYF | TRBV7-9 | TRBJ2-7 | NA | NA | NA | NA | NA |
| 3 | DIE | CASSPdiEAFF | TRBV7-9 | TRBJ1-1 | NA | NA | NA | NA | NA |
| 3 | DIE | CASSadiEQFF | TRBV7-9 | TRBJ2-1 | NA | NA | NA | NA | NA |
| 3 | DIE | CASSPdiedFF | TRBV7-9 | TRBJ1-1 | NA | NA | NA | NA | NA |
| 3 | DIE | CASSLdiEAFF | TRBV7-9 | TRBJ1-1 | NA | NA | NA | NA | NA |
| 6 | DLNT | CASkrdlNTEAFF | TRBV5-5 | TRBJ1-1 | CAALLGPDKLIF | TRAV1-1 | TRAJ34 | 1310 | Pt-08 |
| 6 | DLNT | CASDRGdlNTGELFF | TRBV19 | TRBJ2-2 | CAVRLTGGGNKLTF | TRAV12-2 | TRAJ10 | 2520 | Pt-08 |
| 6 | DLNT | CASSSdlNTGELFF | TRBV11-2 | TRBJ2-2 | CALYSGAGSYQLTF | TRAV9-2 | TRAJ28 | 2718 | Pt-08 |
| 6 | DLNT | CAItdlNTGELFF | TRBV27 | TRBJ2-2 | NA | NA | NA | NA | NA |
| 6 | DLNT | CAGQdlNTGELFF | TRBV7-2 | TRBJ2-2 | NA | NA | NA | NA | NA |
| 6 | DLNT | CASSDlNTGELFF | TRBV2 | TRBJ2-2 | NA | NA | NA | NA | NA |
| 6 | DLNT | CATedlNTGELFF | TRBV6-5 | TRBJ2-2 | NA | NA | NA | NA | NA |
| 6 | DLNT | CASndlNTGELFF | TRBV7-8 | TRBJ2-2 | NA | NA | NA | NA | NA |

| | | | | | | | | | |
|---|---|---|---|---|---|---|---|---|---|
| 6 | DLNT | CASndlNTGELFF | TRBV2 | TRBJ2-2 | NA | NA | NA | NA | NA |
| 6 | DLNT | CAlgdlNTGELFF | TRBV7-2 | TRBJ2-2 | NA | NA | NA | NA | NA |
| 6 | DLNT | CApgdlNTGELFF | TRBV7-2 | TRBJ2-2 | NA | NA | NA | NA | NA |
| 6 | DLNT | CASqdlNTGELFF | TRBV19 | TRBJ2-2 | NA | NA | NA | NA | NA |
| 6 | DLNT | CASqdlNTGELFF | TRBV10-1 | TRBJ2-2 | NA | NA | NA | NA | NA |
| 6 | DLNT | CAIqdlNTGELFF | TRBV13 | TRBJ2-2 | NA | NA | NA | NA | NA |
| 6 | DLNT | CATtdlNTGELFF | TRBV15 | TRBJ2-2 | NA | NA | NA | NA | NA |
| 6 | DLNT | CASydlNTGELFF | TRBV5-1 | TRBJ2-2 | NA | NA | NA | NA | NA |
| 6 | DLNT | CATlvdlNTGELFF | TRBV12-3 | TRBJ2-2 | NA | NA | NA | NA | NA |
| 6 | DLNT | CSAhgdlNTGELFF | TRBV20-1 | TRBJ2-2 | NA | NA | NA | NA | NA |
| 6 | DLNT | CASTdlNTGELFF | TRBV5-4 | TRBJ2-2 | NA | NA | NA | NA | NA |
| 6 | DLNT | CASTdlNTGELFF | TRBV6-5 | TRBJ2-2 | NA | NA | NA | NA | NA |
| 6 | DLNT | CAardqrdlNTGELFF | TRBV2 | TRBJ2-2 | NA | NA | NA | NA | NA |
| 38 | SPD%E | CASSPdNEQFF | TRBV7-6 | TRBJ2-1 | CAVQAATGRRALTF | TRAV20 | TRAJ5 | 375 | Pt-04 |
| 38 | SPD%E | CASSPdsEQYF | TRBV7-9 | TRBJ2-7 | NA | NA | NA | NA | NA |
| 38 | SPD%E | CASSPdiEQFF | TRBV7-9 | TRBJ2-1 | NA | NA | NA | NA | NA |
| 38 | SPD%E | CASSPdiEQYF | TRBV7-9 | TRBJ2-7 | NA | NA | NA | NA | NA |
| 38 | SPD%E | CASSPdiEAFF | TRBV7-9 | TRBJ1-1 | NA | NA | NA | NA | NA |
| 38 | SPD%E | CASSPdiedFF | TRBV7-9 | TRBJ1-1 | NA | NA | NA | NA | NA |
| 38 | SPD%E | CASSPdgEQYF | TRBV7-9 | TRBJ2-7 | NA | NA | NA | NA | NA |
| 44 | SDLD | CASSDlDRaGANVLTF | TRBV6-4 | TRBJ2-6 | CAVRDKPRLMF | TRAV3 | TRAJ31 | 1849 | Pt-02 |
| 44 | SDLD | CAISdlDRGilETQYF | TRBV10-3 | TRBJ2-5 | CAVKTYGGQNFVF | TRAV8-1 | TRAJ26 | 723 | Pt-04 |

| | | | | | | | | | |
|---|---|---|---|---|---|---|---|---|---|
| 44 | SDLD | CASSDldGGEAFF | TRBV2 | TRBJ1-1 | NA | NA | NA | NA | NA |
| 44 | SDLD | CATSDLdsGELFF | TRBV24-1 | TRBJ2-2 | NA | NA | NA | NA | NA |
| 44 | SDLD | CASSDldnlvAFF | TRBV2 | TRBJ1-1 | NA | NA | NA | NA | NA |
| 44 | SDLD | CASSDldTGELFF | TRBV2 | TRBJ2-2 | NA | NA | NA | NA | NA |
| 48 | EQNT | CASSQeqNTEAFF | TRBV3-1 | TRBJ1-1 | CAVGAPGAGSYQLTF | TRAV8-3 | TRAJ28 | 3180 | Pt-08 |
| 48 | EQNT | CAvLAeqNTGELFF | TRBV10-2 | TRBJ2-2 | NA | NA | NA | NA | NA |
| 48 | EQNT | CSVEeqNTGELFF | TRBV29-1 | TRBJ2-2 | NA | NA | NA | NA | NA |
| 48 | EQNT | CASrsrveqNTGELFF | TRBV2 | TRBJ2-2 | NA | NA | NA | NA | NA |
| 48 | EQNT | CASrGLAeqNTGELFF | TRBV2 | TRBJ2-2 | NA | NA | NA | NA | NA |
| 59 | DEVA | CATSDevavGELFF | TRBV24-1 | TRBJ2-2 | CAARRSQGNLIF | TRAV8-6 | TRAJ42 | 2090 | Pt-08 |
| 59 | DEVA | CASSQdevasGELFF | TRBV4-1 | TRBJ2-2 | CAVSPAALGFGNVLHC | TRAV3 | TRAJ35 | 2652 | Pt-08 |
| 59 | DEVA | CSARdevahNTGELFF | TRBV20-1 | TRBJ2-2 | NA | NA | NA | NA | NA |
| 61 | SLD%E | CASSLdwEQYF | TRBV13 | TRBJ2-7 | CIVRHSGGYQKVTF | TRAV26-1 | TRAJ13 | 922 | Pt-02 |
| 61 | SLD%E | CASSLdTEAFF | TRBV12-4 | TRBJ1-1 | CAPGWAGTALIF | TRAV6 | TRAJ15 | 8 | Pt-08 |
| 61 | SLD%E | CASSLdietYF | TRBV7-9 | TRBJ2-7 | NA | NA | NA | NA | NA |
| 61 | SLD%E | CASSLdsEQFF | TRBV7-9 | TRBJ2-1 | NA | NA | NA | NA | NA |
| 61 | SLD%E | CASSLdvEQYF | TRBV7-9 | TRBJ2-7 | NA | NA | NA | NA | NA |
| 61 | SLD%E | CASSLdiEAFF | TRBV7-9 | TRBJ1-1 | NA | NA | NA | NA | NA |
| 79 | S%ANTGE | CASSYaNTGELFF | TRBV5-4 | TRBJ2-2 | CAVGVDSNYQLIW | TRAV39 | TRAJ33 | 1104 | Pt-04 |
| 79 | S%ANTGE | CASSEaNTGELFF | TRBV7-9 | TRBJ2-2 | NA | NA | NA | NA | NA |
| 79 | S%ANTGE | CASSDaNTGELFF | TRBV5-1 | TRBJ2-2 | NA | NA | NA | NA | NA |
| 79 | S%ANTGE | CASSnaNTGELFF | TRBV7-9 | TRBJ2-2 | NA | NA | NA | NA | NA |

| | | | | | | | | | |
|---|---|---|---|---|---|---|---|---|---|
| 82 | S%GTGHQP | CATSkGTGhQPQHF | TRBV15 | TRBJ1-5 | CVVRGSNDYKLSF | TRAV12-1 | TRAJ20 | 827 | Pt-08 |
| 82 | S%GTGHQP | CATSgGTGhQPQHF | TRBV24-1 | TRBJ1-5 | NA | NA | NA | NA | NA |
| 82 | S%GTGHQP | CASSLGTGhQPQHF | TRBV27 | TRBJ1-5 | NA | NA | NA | NA | NA |
| 98 | SIIG | CASSIigNQPQHF | TRBV19 | TRBJ1-5 | CILNSGGSNYKLTF | TRAV26-2 | TRAJ53 | 949 | Pt-02 |
| 98 | SIIG | CASSQsiigGTETQYF | TRBV16 | TRBJ2-5 | CAEKGDKIIF | TRAV5 | TRAJ30 | 547 | Pt-04 |
| 98 | SIIG | CASSIiGGgNTGELFF | TRBV19 | TRBJ2-2 | NA | NA | NA | NA | NA |
| 102 | SKGN | CASSYskgNQPQHF | TRBV6-6 | TRBJ1-5 | CILSRNFGNEKLTF | TRAV26-2 | TRAJ48 | 1621 | Pt-04 |
| 102 | SKGN | CAISEskGNTIYF | TRBV10-3 | TRBJ1-3 | CVVSADNYGQNFVF | TRAV10 | TRAJ26 | 1445 | Pt-08 |
| 102 | SKGN | CSARdskgngwNTGELFF | TRBV20-1 | TRBJ2-2 | NA | NA | NA | NA | NA |
| 166 | SLGTG%D | CASSLGTGdddTF | TRBV7-9 | TRBJ1-2 | CALLGDSNYQLIW | TRAV16 | TRAJ33 | 418 | Pt-08 |
| 166 | SLGTG%D | CASSLGTGGdLHF | TRBV7-9 | TRBJ1-6 | NA | NA | NA | NA | NA |
| 181 | SDGTS%E | CAISdGTSYEQYF | TRBV10-3 | TRBJ2-7 | CAGPGVSSGGSNYKLTF | TRAV25 | TRAJ53 | 3305 | Pt-08 |
| 181 | SDGTS%E | CASSDGTSnEQYF | TRBV5-1 | TRBJ2-7 | NA | NA | NA | NA | NA |
| 192 | SGGTG%QP | CASSgGTGGQPQHF | TRBV2 | TRBJ1-5 | CVVNSDYGQNFVF | TRAV12-1 | TRAJ26 | 1429 | Pt-08 |
| 192 | SGGTG%QP | CATSgGTGhQPQHF | TRBV24-1 | TRBJ1-5 | NA | NA | NA | NA | NA |
| 202 | SFLAGG%TGE | CASSfLAGGqTGELFF | TRBV5-6 | TRBJ2-2 | CAVNKNTGGFKTIF | TRAV8-1 | TRAJ9 | 1869 | Pt-08 |
| 202 | SFLAGG%TGE | CASSfLAGGNTGELFF | TRBV19 | TRBJ2-2 | NA | NA | NA | NA | NA |
| 247 | SD%YG | CASSDdYGYTF | TRBV5-1 | TRBJ1-2 | CAVNAQGNNRLAF | TRAV8-1 | TRAJ7 | 1898 | Pt-02 |
| 247 | SD%YG | CASSDsYGYTF | TRBV7-8 | TRBJ1-2 | NA | NA | NA | NA | NA |
| 259 | SY%NTGE | CASSYaNTGELFF | TRBV5-4 | TRBJ2-2 | CAVGVDSNYQLIW | TRAV39 | TRAJ33 | 1104 | Pt-04 |
| 259 | SY%NTGE | CASSYqNTGELFF | TRBV7-8 | TRBJ2-2 | NA | NA | NA | NA | NA |
| 279 | %YANTGE | CASSYaNTGELFF | TRBV5-4 | TRBJ2-2 | CAVGVDSNYQLIW | TRAV39 | TRAJ33 | 1104 | Pt-04 |

| | | | | | | | | | |
|---|---|---|---|---|---|---|---|---|---|
| 279 | %YANTGE | CAIqyaNTGELFF | TRBV15 | TRBJ2-2 | NA | NA | NA | NA | NA |
| 284 | QQAN | CASSQqaNTEAFF | TRBV3-1 | TRBJ1-1 | CAVTYTGANSKLTF | TRAV8-6 | TRAJ56 | 3280 | Pt-08 |
| 284 | QQAN | CASqqaNTGELFF | TRBV19 | TRBJ2-2 | NA | NA | NA | NA | NA |
| 285 | S%TNTGE | CASSGTNTGELFF | TRBV6-5 | TRBJ2-2 | CAASSSGTYKYIF | TRAV29/DV5 | TRAJ40 | 782 | Pt-02 |
| 285 | S%TNTGE | CASSDtNTGELFF | TRBV5-1 | TRBJ2-2 | NA | NA | NA | NA | NA |
| 285 | S%TNTGE | CASShtNTGELFF | TRBV9 | TRBJ2-2 | NA | NA | NA | NA | NA |
| 288 | DLNS | CAGGgdlnSGNTIYF | TRBV12-3 | TRBJ1-3 | CVGNSGYALNF | TRAV12-1 | TRAJ41 | 480 | Pt-08 |
| 288 | DLNS | CASSDlnSGEQYF | TRBV2 | TRBJ2-7 | NA | NA | NA | NA | NA |
| 288 | DLNS | CATmddlnsGELFF | TRBV2 | TRBJ2-2 | NA | NA | NA | NA | NA |
| 302 | %TENTGE | CAltteNTGELFF | TRBV30 | TRBJ2-2 | CVVNKITGTASKLTF | TRAV12-1 | TRAJ44 | 3701 | Pt-08 |
| 302 | %TENTGE | CATSteNTGELFF | TRBV15 | TRBJ2-2 | NA | NA | NA | NA | NA |
| 385 | SPLAGG%TGE | CASSPLAGGtTGELFF | TRBV13 | TRBJ2-2 | CAMREGGNSGGSNYKLTF | TRAV14/DV4 | TRAJ53 | 1808 | Pt-08 |
| 385 | SPLAGG%TGE | CASSPLAGGNTGELFF | TRBV3-1 | TRBJ2-2 | NA | NA | NA | NA | NA |
| 385 | SPLAGG%TGE | CASSPLAGGNTGELFF | TRBV4-3 | TRBJ2-2 | NA | NA | NA | NA | NA |
| 428 | %GTNTGE | CASSGTNTGELFF | TRBV6-5 | TRBJ2-2 | CAASSSGTYKYIF | TRAV29/DV5 | TRAJ40 | 782 | Pt-02 |
| 428 | %GTNTGE | CASGGTNTGELFF | TRBV2 | TRBJ2-2 | NA | NA | NA | NA | NA |
| 533 | SPDS% | CASSPdSPLHF | TRBV6-6 | TRBJ1-6 | CALSEARNFNKFYF | TRAV19 | TRAJ21 | 3176 | Pt-08 |
| 533 | SPDS% | CASSPdsEQYF | TRBV7-9 | TRBJ2-7 | NA | NA | NA | NA | NA |
| 540 | S%GTSNE | CASSfGTSNEQFF | TRBV27 | TRBJ2-1 | CAETLAGGSTLGRLYF | TRAV5 | TRAJ18 | 448 | Pt-08 |
| 540 | S%GTSNE | CASSDGTSnEQYF | TRBV5-1 | TRBJ2-7 | NA | NA | NA | NA | NA |
| 580 | SL%VE | CSgslgvEQYF | TRBV20-1 | TRBJ2-7 | CAVNFYNQGGKLIF | TRAV8-1 | TRAJ23 | 610 | Pt-08 |
| 580 | SL%VE | CASSLdvEQYF | TRBV7-9 | TRBJ2-7 | NA | NA | NA | NA | NA |

| | | | | | | | | | |
|---|---|---|---|---|---|---|---|---|---|
| 618 | RD%DT | CASrdaDTQYF | TRBV5-5 | TRBJ2-3 | CVVNTGFQKLVF | TRAV12-1 | TRAJ8 | 2922 | Pt-08 |
| 618 | RD%DT | CANrdTDTQYF | TRBV3-1 | TRBJ2-3 | NA | NA | NA | NA | NA |
| 664 | SG%NTGE | CASSGTNTGELFF | TRBV6-5 | TRBJ2-2 | CAASSSGTYKYIF | TRAV29/DV5 | TRAJ40 | 782 | Pt-02 |
| 664 | SG%NTGE | CASSGLNTGELFF | TRBV5-1 | TRBJ2-2 | NA | NA | NA | NA | NA |

**Supplementary Table 4.** GLIPH2 analysis of patient TCRs and known HLA-A*02:01-restricted CMV- and EBV-specific TCRs

| Specificity group | Motif | CDR3b | TRBV | TRBJ | CDR3a | TRAV | TRAJ | Patient |
|---|---|---|---|---|---|---|---|---|
| 15 | SSA%YG | CASSSASYGYTF | TRBV19 | TRBJ1-2 | NA | NA | NA | NLVPMVATV_pp65_CMV |
| 15 | SSA%YG | CASSSANYGYTF | TRBV12-4 | TRBJ1-2 | CAGPMKTSYDKVIF | TRAV35 | TRAJ50 | Pt-08 |
| 15 | SSA%YG | CASSSANYGYTF | TRBV12-4 | TRBJ1-2 | NA | NA | NA | NLVPMVATV_pp65_CMV |
| 15 | SSA%YG | CASSSANYGYTF | TRBV19 | TRBJ1-2 | NA | NA | NA | NLVPMVATV_pp65_CMV |
| 15 | SSA%YG | CASSSANYGYTF | TRBV12-3 | TRBJ1-2 | NA | NA | NA | NLVPMVATV_pp65_CMV |
| 15 | SSA%YG | CASSSANYGYTF | TRBV11-1 | TRBJ1-2 | NA | NA | NA | NLVPMVATV_pp65_CMV |
| 15 | SSA%YG | CASSSAYYGYTF | TRBV12-4 | TRBJ1-2 | NA | NA | NA | NLVPMVATV_pp65_CMV |
| 15 | SSA%YG | CASSSATYGYTF | TRBV19 | TRBJ1-2 | NA | NA | NA | NLVPMVATV_pp65_CMV |
| 15 | SSA%YG | CASSSAHYGYTF | TRBV11-1 | TRBJ1-2 | NA | NA | NA | NLVPMVATV_pp65_CMV |
| 15 | SSA%YG | CASSSAFYGYTF | TRBV19 | TRBJ1-2 | NA | NA | NA | NLVPMVATV_pp65_CMV |
| 15 | SSA%YG | CASSSAFYGYTF | TRBV11-1 | TRBJ1-2 | NA | NA | NA | NLVPMVATV_pp65_CMV |
| 64 | GQAW | CASSSPVQGQAWVYGYTF | TRBV28 | TRBJ1-2 | CAVRPADRGSTLGRLYF | TRAV1-2 | TRAJ18 | Pt-08 |
| 64 | GQAW | CAISDPPGQAWGSPLHF | TRBV10-3 | TRBJ1-6 | CAARYGGATNKLIF | TRAV29/DV5 | TRAJ32 | Pt-08 |
| 64 | GQAW | CASSFGQAWETQYF | TRBV7-9 | TRBJ2-5 | NA | NA | NA | NLVPMVATV_pp65_CMV |
| 64 | GQAW | CASSFGQAWYEQYF | TRBV7-9 | TRBJ2-7 | NA | NA | NA | NLVPMVATV_pp65_CMV |
| 87 | R%GVGNT | CSARTGVGNTIYF | TRBV20-1 | TRBJ1-3 | NA | NA | NA | GLCTLVAML_BMLF1_EBV |
| 87 | R%GVGNT | CSARSGVGNTIYF | TRBV20-1 | TRBJ1-3 | NA | NA | NA | GLCTLVAML_BMLF1_EBV |
| 87 | R%GVGNT | CSARVGVGNTIYF | TRBV20-1 | TRBJ1-3 | NA | NA | NA | GLCTLVAML_BMLF1_EBV |

| | | | | | | | | |
|---|---|---|---|---|---|---|---|---|
| 87 | R%GVGNT | CASREGVGNTIYF | TRBV6-5 | TRBJ1-3 | CAMGGSEKLVF | TRAV29/DV5 | TRAJ57 | Pt-08 |
| 109 | SL%LENE | CASSLGLENEQFF | TRBV7-8 | TRBJ2-1 | NA | NA | NA | GLCTLVAML_BMLF1_EBV |
| 109 | SL%LENE | CASSLTLENEQFF | TRBV7-3 | TRBJ2-1 | CAAKTGANNLFF | TRAV29/DV5 | TRAJ36 | Pt-04 |
| 187 | QRGF | CASRSAQRGFTETQYF | TRBV28 | TRBJ2-5 | NA | NA | NA | NLVPMVATV_pp65_CMV |
| 187 | QRGF | CASSPSTQRGFTAEAFF | TRBV7-2 | TRBJ1-1 | CARNKGGTGFQKLVF | TRAV6 | TRAJ8 | Pt-02 |
| 198 | SFGQA%YE | CASSFGQAWYEQYF | TRBV7-9 | TRBJ2-7 | NA | NA | NA | NLVPMVATV_pp65_CMV |
| 198 | SFGQA%YE | CASSFGQAHYEQYF | TRBV5-1 | TRBJ2-7 | CARLPWRDNYGQNFVF | TRAV12-3 | TRAJ26 | Pt-08 |
| 204 | SLA%GATNEK | CASSLAPGATNEKLFF | TRBV7-6 | TRBJ1-4 | NA | NA | NA | NLVPMVATV_pp65_CMV |
| 204 | SLA%GATNEK | CASSLAPGATNEKLFF | TRBV11-1 | TRBJ1-4 | NA | NA | NA | NLVPMVATV_pp65_CMV |
| 204 | SLA%GATNEK | CASSLAPGATNEKLFF | TRBV7-9 | TRBJ1-4 | NA | NA | NA | NLVPMVATV_pp65_CMV |
| 204 | SLA%GATNEK | CASSLAGGATNEKLFF | TRBV5-1 | TRBJ1-4 | CAAKSGAGSYQLTF | TRAV29/DV5 | TRAJ28 | Pt-04 |
| 225 | SLTTG% | CASSLTTGGYTF | TRBV19 | TRBJ1-2 | CAVGYDMRF | TRAV22 | TRAJ43 | Pt-04 |
| 225 | SLTTG% | CASSLTTGIQYF | TRBV6-2 | TRBJ2-4 | NA | NA | NA | YVLDHLIVV_BRLF1_EBV |
| 239 | S%LLAGYNE | CASSLLLAGYNEQFF | TRBV11-2 | TRBJ2-1 | NA | NA | NA | NLVPMVATV_pp65_CMV |
| 239 | S%LLAGYNE | CASSQLLAGYNEQFF | TRBV4-3 | TRBJ2-1 | CAASSLSGGSNYKLTF | TRAV23/DV6 | TRAJ53 | Pt-04 |
| 292 | SQ%PGGE | CASSQDPGGEQYF | TRBV4-3 | TRBJ2-7 | CAENGGTSYGKLTF | TRAV13-2 | TRAJ52 | Pt-02 |
| 292 | SQ%PGGE | CASSQSPGGEQYF | TRBV14 | TRBJ2-7 | NA | NA | NA | GLCTLVAML_BMLF1_EBV |
| 346 | SLL%SNQP | CASSLLVSNQPQHF | TRBV2 | TRBJ1-5 | CAIVRDDKIIF | TRAV17 | TRAJ30 | Pt-04 |
| 346 | SLL%SNQP | CASSLLGSNQPQHF | TRBV27 | TRBJ1-5 | CAVLPQGGSEKLVF | TRAV36/DV7 | TRAJ57 | Pt-04 |
| 346 | SLL%SNQP | CASSLLGSNQPQHF | TRBV7-8 | TRBJ1-5 | NA | NA | NA | GLCTLVAML_BMLF1_EBV |
| 372 | SLY%DT | CASSLYPDTQYF | TRBV27 | TRBJ2-3 | CAARYSGGGADGLTF | TRAV13-1 | TRAJ45 | Pt-04 |
| 372 | SLY%DT | CASSLYSDTQYF | TRBV28 | TRBJ2-3 | NA | NA | NA | YVLDHLIVV_BRLF1_EBV |

| | | | | | | | | |
|---|---|---|---|---|---|---|---|---|
| 433 | SPGT%KET | CASSPGTDKETQYF | TRBV5-1 | TRBJ2-5 | CAVNRPGGGNKLTF | TRAV12-2 | TRAJ10 | Pt-08 |
| 433 | SPGT%KET | CASSPGTLKETQYF | TRBV12-3 | TRBJ2-5 | NA | NA | NA | NLVPMVATV_pp65_CMV |
| 441 | S%VGGYE | CASSLVGGYEQYF | TRBV5-4 | TRBJ2-7 | CAIRRGGGTSYGKLTF | TRAV13-1 | TRAJ52 | Pt-08 |
| 441 | S%VGGYE | CASSHVGGYEQYF | TRBV4-1 | TRBJ2-7 | NA | NA | NA | GLCTLVAML_BMLF1_EBV |
| 472 | SL%GSNQP | CASSLQGSNQPQHF | TRBV6-2 | TRBJ1-5 | NA | NA | NA | YVLDHLIVV_BRLF1_EBV |
| 472 | SL%GSNQP | CASSLEGSNQPQHF | TRBV7-8 | TRBJ1-5 | CDNAGNMLTF | TRAV27 | TRAJ39 | Pt-08 |
| 472 | SL%GSNQP | CASSLLGSNQPQHF | TRBV27 | TRBJ1-5 | CAVLPQGGSEKLVF | TRAV36/DV7 | TRAJ57 | Pt-04 |
| 472 | SL%GSNQP | CASSLLGSNQPQHF | TRBV7-8 | TRBJ1-5 | NA | NA | NA | GLCTLVAML_BMLF1_EBV |
| 501 | SLG%ENE | CASSLGLENEQFF | TRBV7-8 | TRBJ2-1 | NA | NA | NA | GLCTLVAML_BMLF1_EBV |
| 501 | SLG%ENE | CASSLGEENEQFF | TRBV7-9 | TRBJ2-1 | CAESIFYNAGNMLTF | TRAV5 | TRAJ39 | Pt-08 |
| 568 | S%QGSNQP | CASSPQGSNQPQHF | TRBV18 | TRBJ1-5 | CAASAGNAGKSTF | TRAV29/DV5 | TRAJ27 | Pt-08 |
| 568 | S%QGSNQP | CASSLQGSNQPQHF | TRBV6-2 | TRBJ1-5 | NA | NA | NA | YVLDHLIVV_BRLF1_EBV |
| 592 | S%TGTYE | CATSWTGTYEQYF | TRBV15 | TRBJ2-7 | NA | NA | NA | YVLDHLIVV_BRLF1_EBV |
| 592 | S%TGTYE | CASSLTGTYEQYF | TRBV11-2 | TRBJ2-7 | CAERMAGNMLTF | TRAV13-2 | TRAJ39 | Pt-04 |
| 592 | S%TGTYE | CASSLTGTYEQYF | TRBV4-1 | TRBJ2-7 | CVILTGNQFYF | TRAV29/DV5 | TRAJ49 | Pt-08 |
| 595 | S%GQQET | CASSYGQQETQYF | TRBV6-5 | TRBJ2-5 | CAVRAPGGKLIF | TRAV1-1 | TRAJ23 | Pt-08 |
| 595 | S%GQQET | CASSFGQQETQYF | TRBV12-4 | TRBJ2-5 | NA | NA | NA | GLCTLVAML_BMLF1_EBV |
| 600 | SG%GGTDT | CASSGVGGTDTQYF | TRBV9 | TRBJ2-3 | NA | NA | NA | NLVPMVATV_pp65_CMV |
| 600 | SG%GGTDT | CATSGQGGTDTQYF | TRBV24-1 | TRBJ2-3 | CAGDDAGGTSYGKLTF | TRAV27 | TRAJ52 | Pt-02 |
| 647 | SRGA%E | CAWSRGAVEQFF | TRBV30 | TRBJ2-1 | NA | NA | NA | GLCTLVAML_BMLF1_EBV |
| 647 | SRGA%E | CASSRGAYEQYF | TRBV14 | TRBJ2-7 | CATYGGATNKLIF | TRAV14/DV4 | TRAJ32 | Pt-04 |
| 648 | SL%GYTE | CASSLGGYTEAFF | TRBV7-2 | TRBJ1-1 | CALSGRSGGYQKVTF | TRAV16 | TRAJ13 | Pt-02 |

| | | | | | | | | |
|---|---|---|---|---|---|---|---|---|
| 648 | SL%GYTE | CASSLEGYTEAFF | TRBV11-2 | TRBJ1-1 | NA | NA | NA | NLVPMVATV_pp65_CMV |
| 648 | SL%GYTE | CASSLEGYTEAFF | TRBV27 | TRBJ1-1 | NA | NA | NA | NLVPMVATV_pp65_CMV |
| 680 | S%ANYG | CASSFANYGYTF | TRBV19 | TRBJ1-2 | NA | NA | NA | NLVPMVATV_pp65_CMV |
| 680 | S%ANYG | CASSSANYGYTF | TRBV12-4 | TRBJ1-2 | CAGPMKTSYDKVIF | TRAV35 | TRAJ50 | Pt-08 |
| 680 | S%ANYG | CASSSANYGYTF | TRBV12-4 | TRBJ1-2 | NA | NA | NA | NLVPMVATV_pp65_CMV |
| 680 | S%ANYG | CASSSANYGYTF | TRBV19 | TRBJ1-2 | NA | NA | NA | NLVPMVATV_pp65_CMV |
| 680 | S%ANYG | CASSSANYGYTF | TRBV12-3 | TRBJ1-2 | NA | NA | NA | NLVPMVATV_pp65_CMV |
| 680 | S%ANYG | CASSSANYGYTF | TRBV11-1 | TRBJ1-2 | NA | NA | NA | NLVPMVATV_pp65_CMV |
| 689 | S%GNTE | CASSLGNTEAFF | TRBV12-3 | TRBJ1-1 | CAVREGYSTLTF | TRAV1-1 | TRAJ11 | Pt-08 |
| 689 | S%GNTE | CASSLGNTEAFF | TRBV6-2 | TRBJ1-1 | NA | NA | NA | GLCTLVAML_BMLF1_EBV |
| 689 | S%GNTE | CASSRGNTEAFF | TRBV12-4 | TRBJ1-1 | CILRDSGYSTLTF | TRAV26-2 | TRAJ11 | Pt-08 |
| 689 | S%GNTE | CASSDGNTEAFF | TRBV11-3 | TRBJ1-1 | CAVRTGNQFYF | TRAV8-3 | TRAJ49 | Pt-02 |
| 708 | S%SDT | CASSLSDTQYF | TRBV5-1 | TRBJ2-3 | CALSTQAAGNKLTF | TRAV9-2 | TRAJ17 | Pt-08 |
| 708 | S%SDT | CASSTSDTQYF | TRBV25-1 | TRBJ2-3 | NA | NA | NA | GLCTLVAML_BMLF1_EBV |

**Supplementary Table 5.** Primer sequences for HLA class I typing

| Primer Name | Restriction site (HindIII or BamHI) | Index | Target specific sequence | Final sequence (5' - 3') |
|---|---|---|---|---|
| PCR1_fwd-1 | TACGCCAAGCTT | ACGAGTGCGT | TGGCCCTGACCSAGACCTG | TACGCCAAGCTTACGAGTGCGTTGGCCCTGACCSAGACCTG |
| PCR1_fwd-2 | TACGCCAAGCTT | ACGCTCGACA | TGGCCCTGACCSAGACCTG | TACGCCAAGCTTACGCTCGACATGGCCCTGACCSAGACCTG |
| PCR1_fwd-3 | TACGCCAAGCTT | AGACGCACTC | TGGCCCTGACCSAGACCTG | TACGCCAAGCTTAGACGCACTCTGGCCCTGACCSAGACCTG |
| PCR1_fwd-4 | TACGCCAAGCTT | AGCACTGTAG | TGGCCCTGACCSAGACCTG | TACGCCAAGCTTAGCACTGTAGTGGCCCTGACCSAGACCTG |
| PCR1_fwd-5 | TACGCCAAGCTT | ATCAGACACG | TGGCCCTGACCSAGACCTG | TACGCCAAGCTTATCAGACACGTGGCCCTGACCSAGACCTG |
| PCR1_fwd-6 | TACGCCAAGCTT | ATATCGCGAG | TGGCCCTGACCSAGACCTG | TACGCCAAGCTTATATCGCGAGTGGCCCTGACCSAGACCTG |
| PCR1_fwd-7 | TACGCCAAGCTT | CGTGTCTCTA | TGGCCCTGACCSAGACCTG | TACGCCAAGCTTCGTGTCTCTATGGCCCTGACCSAGACCTG |
| PCR1_fwd-8 | TACGCCAAGCTT | CTCGCGTGTC | TGGCCCTGACCSAGACCTG | TACGCCAAGCTTCTCGCGTGTCTGGCCCTGACCSAGACCTG |
| PCR1_rev | ACCCGGGGATCC | - | GKCCTCGCTCTGGTTGTAGT | ACCCGGGGATCCGKCCTCGCTCTGGTTGTAGT |
| PCR2_fwd-1 | TACGCCAAGCTT | TAGTATCAGC | ACTACAACCAGAGCGAGGMC | TACGCCAAGCTTTAGTATCAGCACTACAACCAGAGCGAGGMC |
| PCR2_fwd-2 | TACGCCAAGCTT | TCTCTATGCG | ACTACAACCAGAGCGAGGMC | TACGCCAAGCTTTCTCTATGCGACTACAACCAGAGCGAGGMC |
| PCR2_fwd-3 | TACGCCAAGCTT | TGATACGTCT | ACTACAACCAGAGCGAGGMC | TACGCCAAGCTTTGATACGTCTACTACAACCAGAGCGAGGMC |
| PCR2_fwd-4 | TACGCCAAGCTT | TACTGAGCTA | ACTACAACCAGAGCGAGGMC | TACGCCAAGCTTTACTGAGCTAACTACAACCAGAGCGAGGMC |
| PCR2_fwd-5 | TACGCCAAGCTT | CATAGTAGTG | ACTACAACCAGAGCGAGGMC | TACGCCAAGCTTCATAGTAGTGACTACAACCAGAGCGAGGMC |
| PCR2_fwd-6 | TACGCCAAGCTT | CGAGAGATAC | ACTACAACCAGAGCGAGGMC | TACGCCAAGCTTCGAGAGATACACTACAACCAGAGCGAGGMC |
| PCR2_fwd-7 | TACGCCAAGCTT | ATACGACGTA | ACTACAACCAGAGCGAGGMC | TACGCCAAGCTTATACGACGTAACTACAACCAGAGCGAGGMC |
| PCR2_fwd-8 | TACGCCAAGCTT | TCACGTACTA | ACTACAACCAGAGCGAGGMC | TACGCCAAGCTTTCACGTACTAACTACAACCAGAGCGAGGMC |
| PCR2_rev | ACCCGGGGATCC | - | TGCCAGGTCAGTGTGATCTC | ACCCGGGGATCCTGCCAGGTCAGTGTGATCTC |
